# Sensory input drives rapid homeostatic scaling of the axon initial segment in mouse barrel cortex

**DOI:** 10.1101/2020.02.27.968065

**Authors:** Nora Jamann, Dominik Dannehl, Robin Wagener, Corinna Corcelli, Christian Schultz, Jochen Staiger, Maarten H.P. Kole, Maren Engelhardt

## Abstract

The axon initial segment (AIS) is an important axonal microdomain for action potential initiation and implicated in the regulation of neuronal excitability during activity-dependent cortical plasticity. While structural AIS plasticity has been suggested to fine-tune neuronal activity when network states change, whether it acts as a homeostatic regulatory mechanism in behaviorally relevant contexts remains poorly understood. Using an *in vivo* model of the mouse whisker-to-barrel pathway in combination with immunofluorescence, confocal analysis and patch-clamp electrophysiological recordings, we observed bidirectional AIS plasticity. Furthermore, we find that structural and functional AIS remodeling occurs in distinct temporal domains: long-term sensory deprivation elicits an AIS length increase, accompanied with an increase in neuronal excitability, while sensory enrichment results in a rapid AIS shortening, accompanied by a decrease in action potential generation. Our findings highlight a central role of the AIS in the homeostatic regulation of neuronal input-output relations.

## Introduction

The axon initial segment (AIS) serves as the cellular substrate for action potential (AP) initiation in most neurons of the central nervous system (Bender and Trussell, 2009; Coombs et al., 1957; Kole and Stuart, 2012). It is characterized by a periodic nanoscale arrangement of actin and scaffolding proteins, particularly Ankyrin-G (ankG) and βIV-spectrin (D’Este et al., 2015; Jenkins et al., 2015a; Leterrier et al., 2015), which tether voltage-gated ion channels, extracellular membrane proteins and receptors at the axonal membrane (reviewed in (Leterrier, 2018; Rasband, 2010)). In particular the clustering of sodium and potassium channels at the AIS plays an important role in giving rise to the AP initiation ((Hu et al., 2009; Kole et al., 2008; Kole et al., 2007)).

Recent work has shown that the AIS dynamically adapts its anatomical length and/or location within the axon, relative to the soma, thereby modulating neuronal input/output properties ((Chand et al., 2015; Evans et al., 2015; Grubb and Burrone, 2010; Kuba et al., 2010; Wefelmeyer et al., 2015), reviewed in (Jamann et al., 2018; Kole and Brette, 2018)). Depending on the excitation state of the neuronal network both *in vitro* and *in vivo*, individual neurons can undergo structural and/or functional AIS plasticity, including modulation of channel density or composition, to adapt to or strengthen the changes in synaptic drive ((Bender et al., 2010; Benned-Jensen et al., 2016; Lezmy et al., 2017; Muir and Kittler, 2014); reviewed in (Jamann et al., 2018; Petersen et al., 2016)). For example, triggering chronic depolarization in neuronal networks *in vitro* (Evans et al., 2015; Grubb and Burrone, 2010) or using a complete and irrevocable elimination of sensory input *in vivo* (Kuba et al., 2010) led to significant geometrical and functional AIS changes. These observations are in good support of AIS remodeling observed in disease or injury models (reviewed in (Buffington and Rasband, 2011). In addition to the up- or downregulation of excitability by AIS changes, cell type-specific remodeling of the AIS has also been found to occur either via long-term (days to weeks) (Gutzmann et al., 2014; Kim et al., 2019; Kuba et al., 2010; Schlüter et al., 2017; Vascak et al., 2017)) or short-term manipulations of a few hours (Bender et al., 2010; Evans et al., 2015; Lezmy et al., 2017; Martinello et al., 2015)).

Such adaptations in the AIS have often been hypothesized to reflect homeostatic mechanisms at the network level to ensure a dynamic equilibrium of neuronal activity (reviewed in (Wefelmeyer et al., 2016). However, such a role of the AIS *in vivo* remains to be demonstrated. To explore AIS plasticity in a behaviorally-relevant context, we utilized the mouse whisker-to-barrel system (Feldmeyer et al., 2013; Stuttgen and Schwarz, 2018). This pathway conveys tactile information from vibrissae on the rodent whisker pad to the barrel field in primary somatosensory cortex (S1BF) in a strictly organized hierarchy and is one of the best-studied sensory systems (reviewed in (Petersen, 2007; Wu et al., 2011)). Thalamic input is largely received by layer IV spiny stellate neurons and propagated to supragranular (layer II/III) neurons; infragranular (layer V) principal neurons also receive direct thalamic input (Sermet et al., 2019). S1BF undergoes activity-dependent maturation with several critical periods during postnatal development (reviewed in (Erzurumlu and Gaspar, 2012; Jamann et al., 2018)), ultimately resulting in a precise somatotopic representation of each whisker in a cortical barrel layer IV (Feldmeyer et al., 2013).

Here, we employed a range of sensory deprivation and enrichment paradigms in behaving mice. We investigated whether the AIS geometry and AP generation were affected following these modifications of sensory input, and which time-course these changes followed. We found that AIS plasticity follows two distinct temporal patterns: deprivation of the whisker-to-barrel pathway for 15 days or longer lead to long-term AIS elongation in S1BF layer II/III neurons, accompanied by an increase in pyramidal neuron excitability. In contrast, increasing the activity of the same neuronal population for just 1-3 hours (h) via exposure to an enriched environment resulted in AIS shortening and increased the threshold for AP generation, reducing pyramidal neuron output. In summary, our results indicate a temporally diverse, bidirectional, activity-dependent remodeling of the AIS and input-output properties, supporting its role for homeostatic adaptation under physiological conditions *in vivo*.

## Results

### The AIS undergoes periods of structural plasticity during development

To examine the time course of AIS development in the whisker-to-barrel system, we first investigated normal AIS maturation at selected time points during S1BF development of mice. Based on immunofluorescent staining, AIS length was quantified from the late embryonic period (E20) throughout postnatal development (P1, P3, 7, 13, 15, 21, 28, 35) into adulthood (P45, P180; each time point *n* = 6 animals, at least 100 AIS per animal in layers II/III and V of S1BF; Fig. 1A-B). The AIS scaffolding protein βIV-spectrin, a well-established target for morphometrical analysis of the AIS (Gutzmann et al., 2014; Kim et al., 2019; Schlüter et al., 2017) was utilized, while immunoblot analysis was directed against the known isoforms of ankG, the main regulator of AIS assembly and maintenance (Jenkins et al., 2015b; Rasband, 2011). In line with previous reports (Galiano et al., 2012; Gutzmann et al., 2014), we found that the AIS elongated during the early postnatal period until the end of the second postnatal week, a time at which mice begin active whisking to explore their environment (Arakawa and Erzurumlu, 2015), indicated by grey boxes in Fig.1B). With the onset of active whisking behavior, however, a significant shortening of AIS length was observed both in pyramidal neurons of layers II/III and V (Fig 1B). Adult animals then maintained an intermediate AIS length until P45, after which no more significant length changes were observed (Fig 1B). Although the overall developmental profiles of layers II/III and V were comparable, AIS length in layer V started to decrease after P10 (Fig. 1B). AIS in layer V were generally shorter than in layer II/III (average adult S1BF layer II/III: ∼22 µm, layer V: ∼19 µm) in accordance with previously published data from visual and somatosensory cortex (Gutzmann et al., 2014; Vascak et al., 2017).

**Fig. 1:**
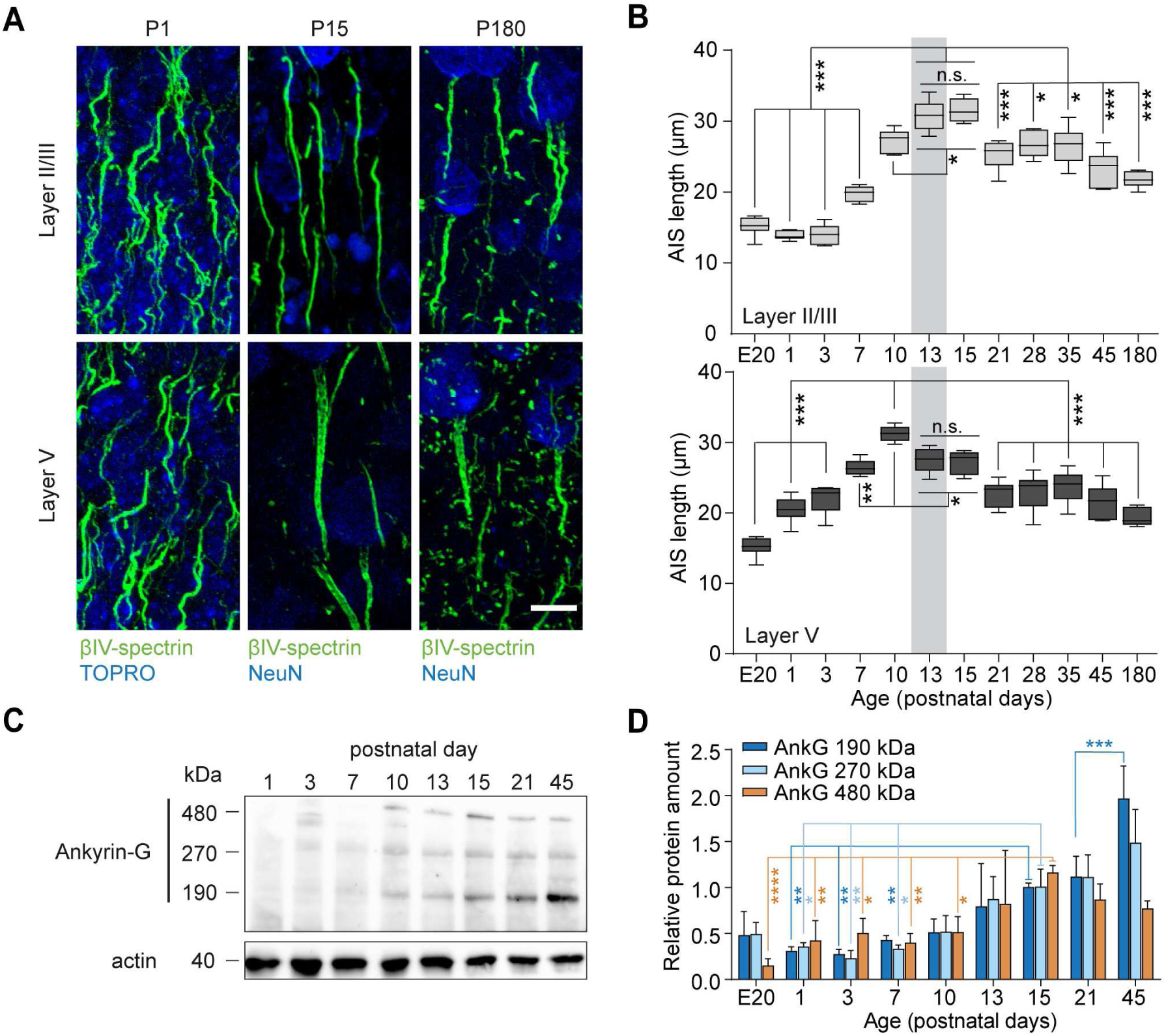
AIS development in S1BF *in vivo*. **A** Representative confocal images of AIS length maturation for three postnatal ages (P1, P15, and P180) in cortical layers II/III and V. Immunostaining against ßIV-spectrin (green), TOPRO (blue) or NeuN (blue) as indicated. Note the long AIS at P15 in both layers. Scale bar 10 µm. **B** Population data of AIS lengths from E20 to P180 in layer II/III (*top*) and V (*bottom*). Initially, AIS length increases until P13-15, after which it decreases. Gray bars indicate the onset of active exploration and whisking (P12–14). Adult animals maintain an intermediate AIS length throughout life (for layer II/III and layer V, One-way ANOVA *P* < 0.001, Holm-Sidak’s post-hoc comparisons, >100 AIS per animal, *n* = 5-6 mice per age group; for all comparisons, \**P* < 0.05, \*\**P* < 0.01, \*\*\**P* < 0.001). Boxplots indicate median with 25 to 75% interval and error bars show min. to max. values. **C** Representative immunoblot of the major ankG isoforms (190, 270, 480 kDa) for postnatal developmental stages. Actin was used as a loading control. **D** Quantification of immunoblot data. Protein expression of the main ankG 480 kDa isoform peaks at P15 (Two-way ANOVA *P* < 0.0001 for time, *P* = 0.07 for isoform, *P* = 0.0007 for the interaction, *n* = 3 mice per age group, Tukey’s multiple comparisons tests \**P* < 0.05, \*\**P* < 0.01, \*\*\**P* < 0.001, only selected statistical comparisons depicted for better visualization of data). Data shown as mean ± SD.

Previous studies have shown that the giant (480 kDa) isoform of *Ank3*, which encodes ankG, serves as the major organizer of AIS assembly (Jenkins et al., 2015b). To test whether the developmental increase in AIS length is accompanied by an upregulation of the ankG isoforms (190, 270 and 480 kDa), we performed immunoblot analysis (Fig. 1C, D). The results showed that consistent with the AIS elongation during the early postnatal period (P1 to P15), there was an increase in all ankG isoforms (Fig. 1D). Interestingly, the expression of the 190 kDa ankG isoform further increased until adulthood, which may be in part explained by synaptogenesis and targeting of this isoform to postsynaptic structures (Tseng et al., 2015).

### AIS elongation is triggered by long-term sensory deprivation

The significant reduction in AIS length seen from P15 (in layer II/III) and P10 (in layer V) onwards coincides with the onset of explorative, active whisking behavior in mice (Arakawa and Erzurumlu, 2015). A similar AIS maturation pattern (early postnatal elongation, followed by length reduction and subsequent stable adult length) was previously described for the primary visual cortex and shown to be regulated by the onset of vision at P13-14 (Gutzmann et al., 2014). We therefore hypothesized that the structural remodeling of the AIS occurs due to developmental changes in tactile input when mice begin to actively explore their environment. We tested this hypothesis by conducting sensory deprivation experiments via daily, bilateral whisker trimming from birth (P0) to P15, P21, and P45, respectively (Group 1, Fig. 2A–C). P15 coincides with the closure of the critical period for layer IV to II/III synapses (Erzurumlu and Gaspar, 2012; Maravall et al., 2004), while P21 was identified as the time point of significant AIS length reduction during development (Fig. 1B). At the P15, P21, and P45 endpoints, mice were perfused and S1BF cryosections examined via immunofluorescence and morphometrical analysis of AIS length in layer II/III (Fig. 2B) and layer V (Fig. S1A) neurons. Bilateral whisker trimming resulted in a significant lengthening of AIS at all time points in Group 1 in layers II/III (Two-way ANOVA *P* < 0.0001 (deprivation), Fig. 2B, C), but not in layer V (Two-way ANOVA *P* = 0.20 (deprivation), Fig. S1A). In a second group, we applied a shorter trimming period of 5 days (P10 to P15). However, this did not lead to AIS length changes in layer II/III (Two-way ANOVA P = 0.3267 (deprivation) Fig. 2A, D), indicating a minimum of 2 weeks of deprivation to induce significant length changes during development. Length frequency distribution analysis of the AIS in Group 1 showed that with whisker trimming until P15, the number of longer AIS increased (Kolmogorov-Smirnov test *P* < 0.0001), with some AIS reaching even 60 µm (Fig. 2E). Therefore, the width of the curve was more than twice as wide in the deprived group (full width at half maximum (FWHM) P15 Ctrl 11.9 µm vs P15 Dep 25.3 µm). By contrast, trimming until P21 led to a shift in the length median, but the full width at half maximum remained similar to control AIS (Kolmogorov-Smirnov test P < 0.0001, FWHM P21 Ctrl 9.79 µm vs P21 Dep 10.86 µm, respectively, Fig. 2E).

**Fig. 2:**
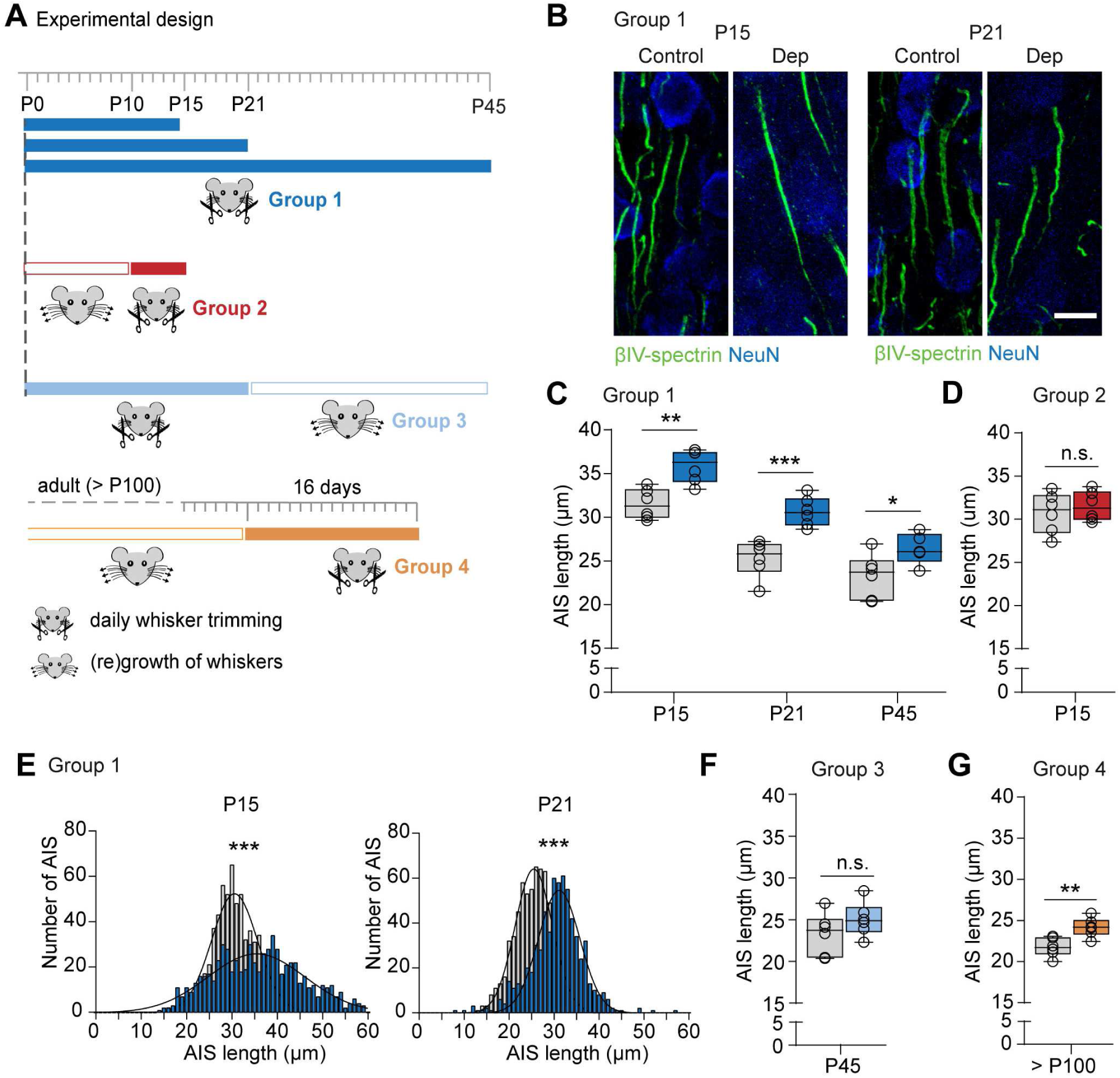
AIS elongation in layer II/III occurs after long-term sensory deprivation in young and adult mice. **A** Schematic of the experimental designs for whisker trimming. Group 1, daily bilateral trimming from P0 to P15, P21 or P45, respectively. Group 2, daily bilateral trimming from P10 to P15. Group 3, daily bilateral trimming until P21, followed by regrowth until P45. Group 4, daily bilateral trimming for 16 days in adult mice (> P100). **B** Representative confocal images of layer II/III AIS stained for ßIV-spectrin (green) and NeuN (blue) in control and deprived mice. Scale bar 10 µm. **C** Population data for Group 1 for control (grey) and deprived (dark blue) animals. Deprivation leads to longer AIS (Two-way ANOVA for deprivation and development *P* < 0.0001, for the interaction *P* = 0.417, >100 AIS/animal, *n* = 5-6 mice, Sidak’s multiple comparison tests \**P* = 0.0416, \*\**P* = 0.002, \*\*\**P* = 0.0002,). **D** Population data for Group 2 for control (grey) and deprived (red) animals. Deprivation from P10 – P15 did not lead to significant length changes. (Two-way ANOVA P = 0.327 for deprivation, P < 0.0001 for age, P = 0.326 for the interaction, Sidak’s multiple comparisons P > 0.05 for all comparisons, n = 6) **E** AIS length frequency histograms for the P15 and P21 time points for control (grey) and deprived (dark blue) animals (for P15 and P21 Dep vs Ctrl, >600 AIS per condition, Kolmogorov-Smirnov test *P* < 0.0001, full width at half maximum P15 Ctrl 11.9 µm vs P15 Dep 25.3 µm; full width at half maximum P21 Ctrl 9.8 µm vs P21 Dep 10.9 µm). **F** Population data for Group 3 for deprived (light blue) and age-matched control animals (gray) at P45. If whiskers were allowed to regrow, AIS length returned to control levels at P45 (unpaired *t*-test *P* = 0.219, >100 AIS/animal, *n* = 6 mice). **G** Population data for adult animals (group 4, > P100) for deprived animals (orange) and age-matched controls (grey). After two weeks of whisker trimming, deprived animals showed significantly longer AIS in layer II/III (unpaired *t*-test \*\**P* = 0.0046, >100 AIS/animal, *n* = 6 mice). Boxplots indicate median with 25 to 75% interval and error bars show minimum to maximum values.

To test whether AIS can recover from deprivation-induced lengthening, another group of mice (Group 3) was trimmed daily from P0 to P21 and sacrificed at P45, after whiskers had regrown (Fig. 2A, F). The results showed that in this group, AIS length returned to mature levels (unpaired *t*-test *P* = 0.22, Fig. 2F). Again, we observed no length differences in layer V neurons (unpaired *t*-test *P* = 0.879 Fig. S1B). To control for any influence of sensory stimulation by the handling or trimming itself, we performed additional control experiments in which animals where handled daily and whiskers were ruffled, but not trimmed. In these animals, no changes in AIS length were observed (unpaired *t*-test *P* = 0.156 Fig. S1C). Finally, to test whether AIS plasticity would also occur in adult mice, we applied whisker trimming after P100 for 16 days (Group 4). Strikingly, a significant AIS elongation was still observed in layer II/III (unpaired *t*-test, *P* = 0.0046; Fig. 2G), but not in layer V (unpaired *t*-test *P* = 0.95, Fig. S1D).

In summary, these experiments indicate that an AIS length increase is induced by long-term sensory deprivation in the whisker-to-barrel system also in mature cortices, and can be reversed by restoration of tactile input.

### Long-term deprivation leads to changes in cellular excitability

In theory, a longer AIS should correspond to increased neuronal excitability by reducing the threshold for AP generation (Kole and Brette, 2018; Kuba et al., 2010). In order to examine whether sensory-deprivation induced AIS length plasticity was associated with electrophysiological changes, we performed whole-cell patch clamp recordings in control and deprived mice, trimmed daily bilaterally from P0 to P15. At P15 (± 2 days), acute slices were prepared and we measured the active and passive properties of layer II/III pyramidal neurons (Dep, *n* = 18 cells from 7 mice; Ctrl, *n* = 15 cells, 7 mice). The data showed that resting membrane properties (incl. resting membrane potential and input resistance) were not changed between the deprivation and control group (Table S1). To assess AP threshold and firing properties, we injected depolarizing current steps (Fig. 3A, C). The results indicated that deprivation significantly increased firing rates near threshold (Two-way ANOVA *P* < 0.0001 for the factor deprivation, Holm-Sidak’s multiple comparisons, for 100 pA, *P* = 0.0037 and for 150 pA, *P* = 0.0135). However, deprivation did not affect the maximum firing rates (∼20 Hz). Furthermore, the current at the maximum slope was significantly reduced in the deprivation group (unpaired *t*-test *P* = 0.028, Fig. 3B). Since the resting membrane properties were not different between the groups (Table S1), these results suggest that the threshold for APs may be reduced. Using 20 ms duration step current injections, we quantified threshold properties for AP generation and found that deprivation led to a ∼50 pA reduction in current threshold, without changing the voltage threshold (unpaired *t*-tests, *P* = 0.015 and *P* = 0.07, respectively; Fig. 3C). Furthermore, neither AP half-width, amplitude nor phase-plane properties of the AP were different between the two groups (Fig. S2A, B). The reduced current threshold for AP generation is consistent with the increased AIS length identified in the population analysis (Fig. 2). In order to test such a link more directly, we performed correlation analyses between the current threshold and the specific AIS length, by immunostaining post-hoc against βIV-spectrin and the biocytin fill of the recorded neuron (Fig. 3D). In agreement with our hypothesis, these data revealed a significant correlation (*r*^2^ = 0.36, *P* = 0.029). Finally, we characterized the properties of synaptic input onto layer II/III neurons by measured spontaneous postsynaptic currents (PSCs) at –70 mV. The deprived cells received PSCs at a higher frequency, albeit with on average a lower amplitude (Fig. 2E, unpaired *t*-test *P* = 0.004 and *P* = 0.0353, respectively).

**Fig. 3:**
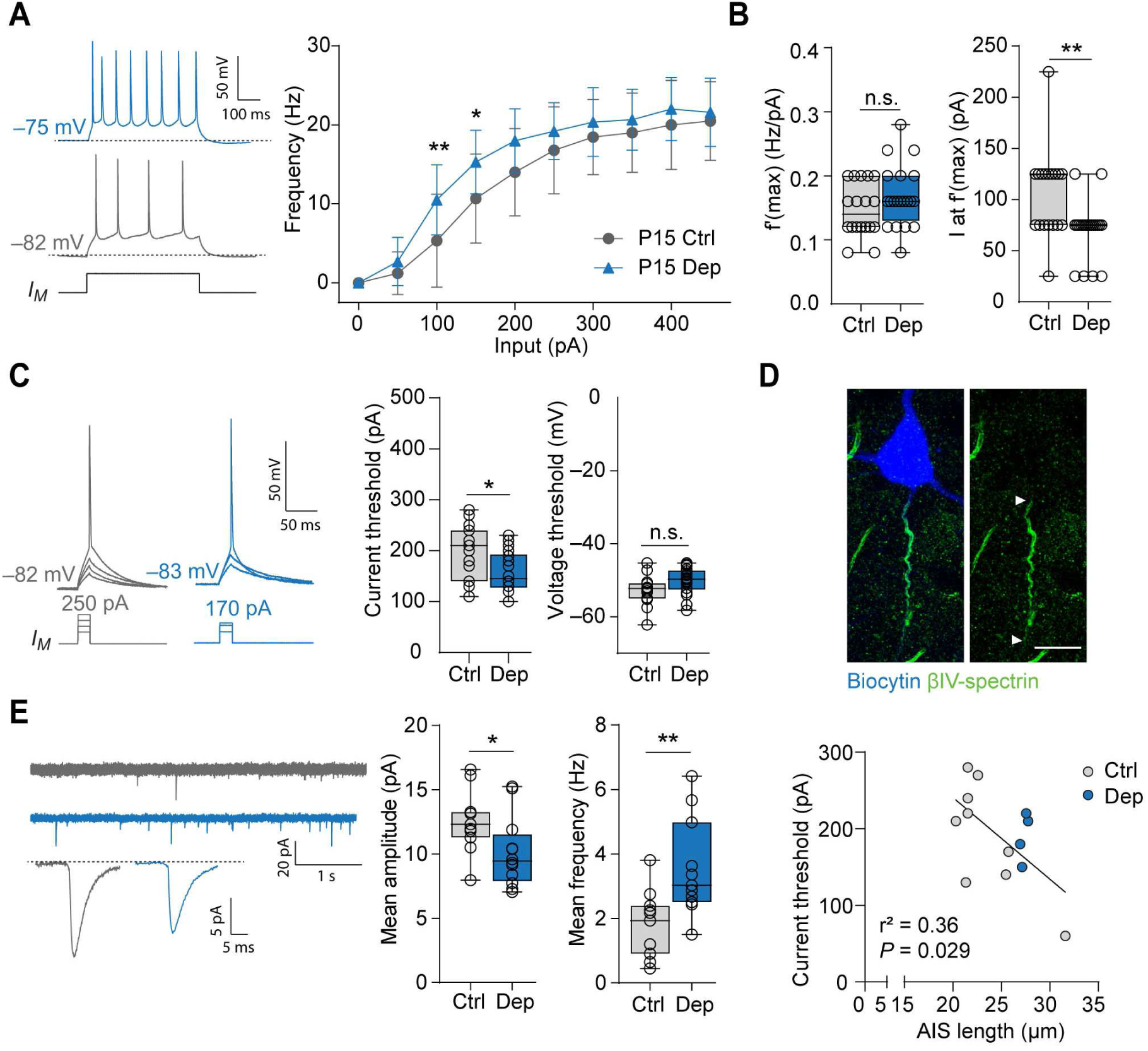
Whisker trimming increases neuronal excitability in layer II/III pyramidal neurons in S1BF. **A** *Left:* Representative traces of AP trains elicited by 100 pA current injections (500 ms pulse). Note the increased firing frequency in the deprived neurons (blue trace). *Right:* Input/frequency relationship as determined by 500 ms long injections of increasing currents. The deprivation group showed significantly increased firing frequencies (Two-way ANOVA *P* < 0.0001 for current injection and deprivation, Holm-Sidak’s multiple comparison test \**P* = 0.0135, \*\**P* = 0.0037, n = 15 cells for Ctrl, 20 cells for Dep) **B** Analysis of the maximum slope f’(max) of the I-*f* curve and the current at the maximum slope I at f’(max) for the respective neuron. The current at the maximum slope was significantly lower in the deprivation group (Mann-Whitney test \*\**P* = 0.0028). The maximum slope was unchanged (unpaired t-test *P* = 0.15). *n* = 14 cells for Ctrl, 18 cells for Dep **C** *Left*: Representative traces of single APs elicited by 20 ms current injections. Injected current was increased in 10 pA increments to determine the current threshold. *Right*: Analysis of current and voltage threshold for threshold APs. Deprived neurons had significantly lower current threshold, the voltage threshold however was unchanged (unpaired *t*-test, current threshold \**P* = 0.0148, voltage threshold *P* = 0.07 *n* = 15 cells for Ctrl, 18 cells for Dep). **D** *Top*: Representative confocal image of a neuron from the P15 deprivation group that was filled with biocytin (blue) for post-hoc determination of AIS length via labeling with βIV-spectrin (green). Arrows indicate start and end of AIS. Scale bar 10 µm. *Bottom*: Correlation analysis of the relationship between AIS length and current threshold. Results of linear regression analysis indicated in figure (*n* = 13 cells). ***E***: *Left*: Representative traces of spontaneous postsynaptic currents (PSCs) recorded at −70 mV. Top trace shows 10 s of recording. Bottom trace depicts the averaged PSCs across the entire recording session of 2 minutes for two example cells. *Right*: Mean amplitude and frequency of PSCs. Deprived neurons received significantly more inputs, which had on average a lower amplitude (unpaired *t*-test, amplitude: \**P* = 0.035, frequency: *\*\*P* = 0.004, *n* = 11 cells for Dep and Ctrl). Boxplots indicate median with 25 to 75% interval and error bars show minimum to maximum values.

Taken together, these data show that a long-term reduction of sensory tactile input increases the intrinsic excitability of layer II/III pyramidal neurons and correlates with increased AIS length.

### AIS undergo rapid structural plasticity after induced exploratory activity *in vivo*

Our data so far suggest that AIS increases in length require long periods of deprivation (days to weeks). In contrast, studies applying depolarizing conditions *in vitro* showed that AIS length changes can be triggered more rapidly, within minutes to hours of increased *in vitro* network activity (Benned-Jensen et al., 2016; Evans et al., 2015). We next tested whether rapid AIS plasticity could be evoked *in vivo* in a behaviorally relevant context, by placing young adult wild type mice at P28 (n = 5-6 per group) to an enriched environment (EE) and expose whiskers to a larger range of novel stimuli. For EE, we placed a variety of novel objects and different types of bedding in a large home cage, enabling enhanced explorative behavior. 12 h prior to the experiment, mice were subjected to unilateral whisker trimming (Fig. 4A). Consequently, the S1BF contralateral to the intact whisker pad received higher activation via the whisker-to-barrel pathway (EE and whiskers intact side) compared to the ipsilateral side (Ctrl side, (Feldman and Brecht, 2005)). To exclude possible short-term deprivation effects of whisker trimming on AIS length, we included a “0 h” group, where animals that received unilateral whisker trimming were sacrificed immediately after the 12 h post whisker trimming without being placed in the EE cage (Fig. 4A). Experimental groups remained in EE for 1 h, 3 h, and 6 h, respectively. At each end point, mice were sacrificed and S1BF tissue blocks were processed for morphometrical analysis of AIS parameters (n = 5–6 per time point, at least 100 AIS per animal, Fig. 4). Activation of neurons on the EE side of S1BF was confirmed by immunofluorescent detection of the immediate early gene product c-Fos (Fig. 4B), which is upregulated within very short time frames after an increase in neuronal activity occurs (Hughes et al., 1999; Staiger et al., 2002; Wagener et al., 2016). At the 0 h time point, only very scarce immunosignal was detectable (< 1% c-Fos-positive (c-Fos^+^) of all analyzed AIS-endowed cells in layer II/III of S1BF; Fig. 4B, C). In contrast, 1 h after EE exposure, layer II/III neurons showed a significant upregulation in c-Fos expression (∼19% c-Fos^+^; Fig. 4B, C), while control cells remained mostly negative for c-Fos (∼2% c-Fos^+^, Fig. 4B, C). c-Fos was further increased in expression after 3 h (∼38% EE vs. ∼6% Ctrl, Fig. 4B, C) and decreased to baseline levels after 6 h of EE exposure (Fig. 4B, C).

**Fig. 4:**
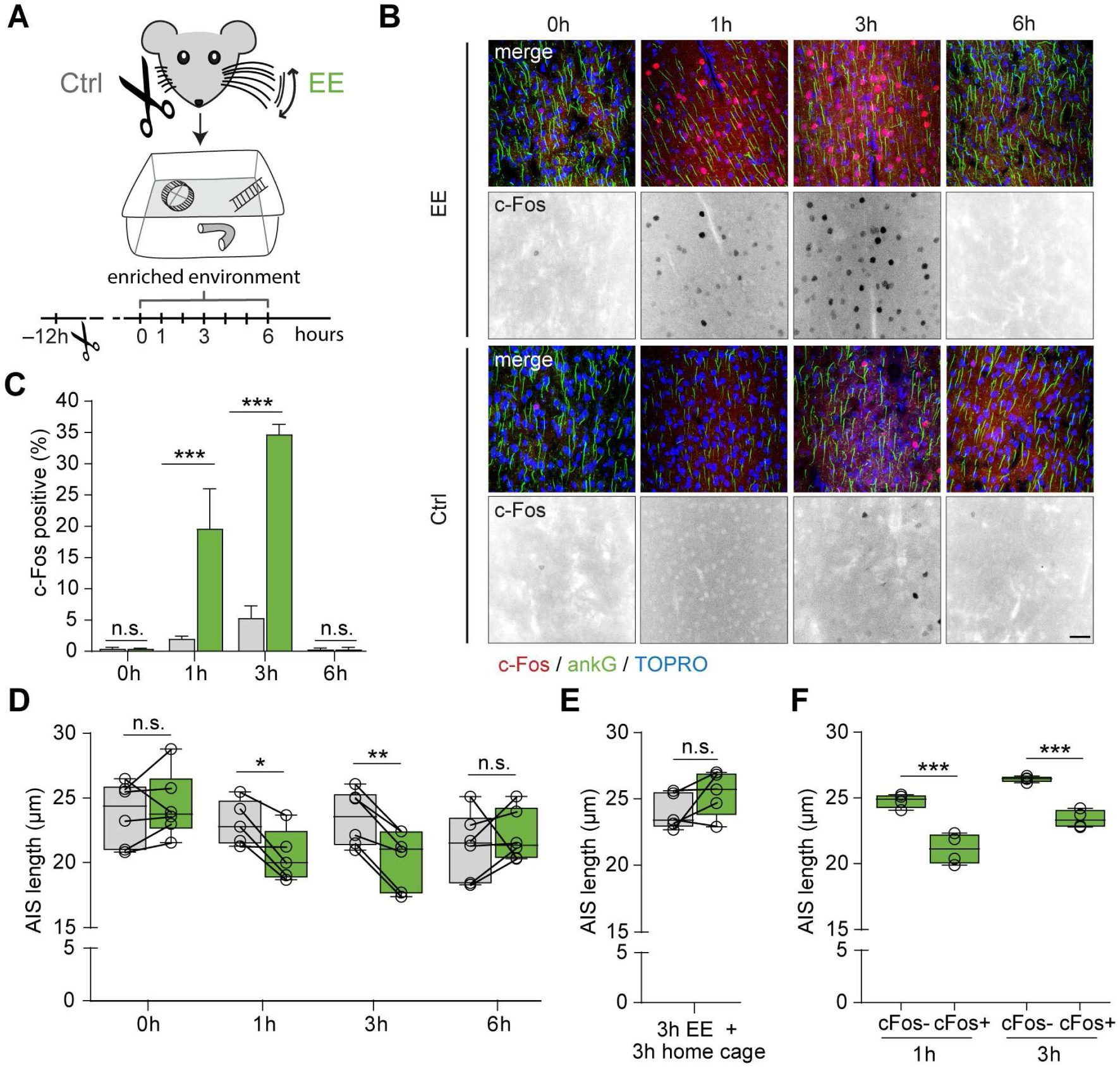
AIS in layer II/III undergo rapid structural remodeling after exposure to an enriched environment (EE) **A** Experimental design: P28 mice (*n* = 5-6 per time point) were exposed to EE conditions 12 h after unilateral whisker trimming. Mice remained in EE for 0 h, 1 h, 3 h, and 6 h, respectively, and were sacrificed for immunofluorescent staining and morphometrical analysis immediately after this period. Layer II/III S1BF pyramidal neurons contralateral to the remaining whiskers represented the stimulated side (EE), while ipsilateral S1BF served as control (Ctrl). **B** Representative confocal images of EE and Ctrl S1BF after immunostainings of layer II/III principal neurons at 1, 3, and 6 h EE exposure. AIS demarked by staining against AnkG (green) with TOPRO as nuclear stain (blue). Staining against the immediate early gene product c-Fos (red) served as indicator of neuronal activity. Inverted black & white panels highlight the increasing c-Fos signal over time. Scale bar 30 µm. **C** Quantification of c-Fos expression indicates rapid and significant upregulation of the marker after 1 h of EE (19.6% c-Fos^+^ neurons) with a peak expression 3 h after EE (38% c-Fos^+^ neurons), and subsequent downregulation 6 h after EE (1% c-Fos^+^ neurons). Two-way ANOVA (EE x time) *P* < 0.0001 for time, EE and interaction of the two factors, Sidak’s multiple comparisons test \*\*\**P* < 0.0001, *n* = 3 animals). **D** Quantification of AIS length reveals a significant length reduction in S1BF layer II/III pyramidal neurons after 1 h and 3 h of EE. AIS length reduction is normalized 6 h after EE induction (Two-way ANOVA (time x EE) *P* = 0.02 for the factor EE, *P* = 0.156 for the factor time (ns), *P* = 0.005 for the interaction of the two, Sidak’s multiple comparisons test \**P* = 0.043, \*\**P* = 0.0051, *n* = 5 - 6 animals; > 100 AIS/animal). Lines indicate matching data from the two hemispheres of individual mice. **E** In a separate experiment, mice were first subjected to 3 h of EE and then placed in home cage conditions for another 3 h. In the home cage environment, AIS length normalized to control levels. (unpaired *t*-test P = 0.195, *n* = 5, > 100 AIS/animal). **F** Subpopulation analysis of AIS length of c-Fos^+^ vs c-Fos^-^ cells in the EE hemisphere reveals significantly decreased AIS length in c-Fos^+^ neurons (One-way ANOVA *P* < 0.0001, Sidak’s multiple comparisons test \*\*\**P* < 0.0001). **D – F** Boxplots indicate median with 25 to 75% interval and error bars show minimum to maximum values.

Next, we asked whether these rapid changes in S1BF network activity after EE triggered structural AIS remodeling in layer II/III cortical neurons. Population analysis of mean AIS length in layer II/III in the EE and Ctrl hemispheres revealed an AIS length reduction at 1 h after EE exposure (Fig. 4E, Two-way RM ANOVA, Sidak’s multiple comparisons test *P* < 0.05). This effect was significantly increased 3 h after EE, leading to a shortening of on average ∼3 µm (Fig.4E, Two-way RM ANOVA, Sidak’s multiple comparisons test *P* < 0.01). AIS length subsequently returned to control levels after 6 h of EE (Fig. 4E). No significant AIS length alterations were detected in layer V (Fig. S3A).

To test whether the return to baseline after 6 h was associated with a reduction in exploratory behavior and thus sensory input, we examined a group of mice which were placed in EE for 3 h and subsequently placed back into their home cage environment for another 3 h. The AIS length in this group remained unchanged in comparison to controls (Fig. 4F, unpaired *t*-test, *P* = 0.195), suggesting a rapid reversibility when sensory input is decreased to home cage environment levels.

Finally, we asked whether the AIS shortening we observed in layer II/III would be selective in neurons activated by the EE exposure (c-Fos^+^ neurons). AIS length of c-Fos^+^ neurons was analyzed by immunostaining against ankG in combination with c-Fos and detection of nuclei via TOPRO in order to identify c-Fos negative (c-Fos^-^) neurons and their corresponding AIS. Consistent with the role of excitability, we found that AIS were significantly shorter (∼3 µm) in c-Fos^+^ neurons compared to surrounding c-Fos^-^ neurons both 1 and 3 h post EE exposure (Fig. 4F, One-way ANOVA *P* < 0.0001).

Taken together, we demonstrate that as little as 1h of exposure to an enriched environment triggers an AIS shortening that requires increased sensory input via the whisker pathway.

### Rapid structural AIS remodeling triggers changes in neuronal excitation

What are the functional consequences of the observed rapid structural AIS changes in pyramidal neurons of S1BF layer II/III? To address this question, we performed whole-cell patch clamp recordings in acute slices in mice that were exposed to 3 h of EE as outlined in Fig. 4A. The passive properties were unchanged between the two groups (Table S2). However, EE neurons generated APs at significantly lower frequencies (mean difference on average 2.4 Hz, Two-way ANOVA, *P* < 0.0001 for the factor EE, Fig. 5A). Consistent with these data, the maximum slope of the I/f curve was significantly lower in EE cells (unpaired *t*-test *P* < 0.05, Fig. 5B). Accordingly, we also detected a significantly higher AP current threshold in EE cells of ∼70 pA (unpaired *t*-test *P* < 0.05, Fig. 5C). Other AP properties such as voltage threshold (Fig. 5C), half-width duration, and amplitude (Fig. S4A) as well as the axonal and somatic components in the phase-plane trajectories of the AP (Fig. S4B) all remained unchanged. To exclude that AIS length and therefore electrophysiological properties would change during incubation of acute slices in ACSF, we plotted the time of the start of the whole-cell recording relative to the time of slice preparation, but found no correlation in current threshold (Fig. S4C). However, consistent with the presumed role of the AIS, post-hoc correlation analysis between AIS length and threshold in biocytin-filled cells revealed that neurons with a higher current threshold also had shorter AIS (Fig. 5D). Since our deprivation experiments indicated changes to synaptic input (Fig. 3E), we recorded PSCs after exposure to EE (Fig. 5E). However, we did not detect any significant differences of mean amplitude or frequency of PSCs (Fig. 5E).

**Fig. 5:**
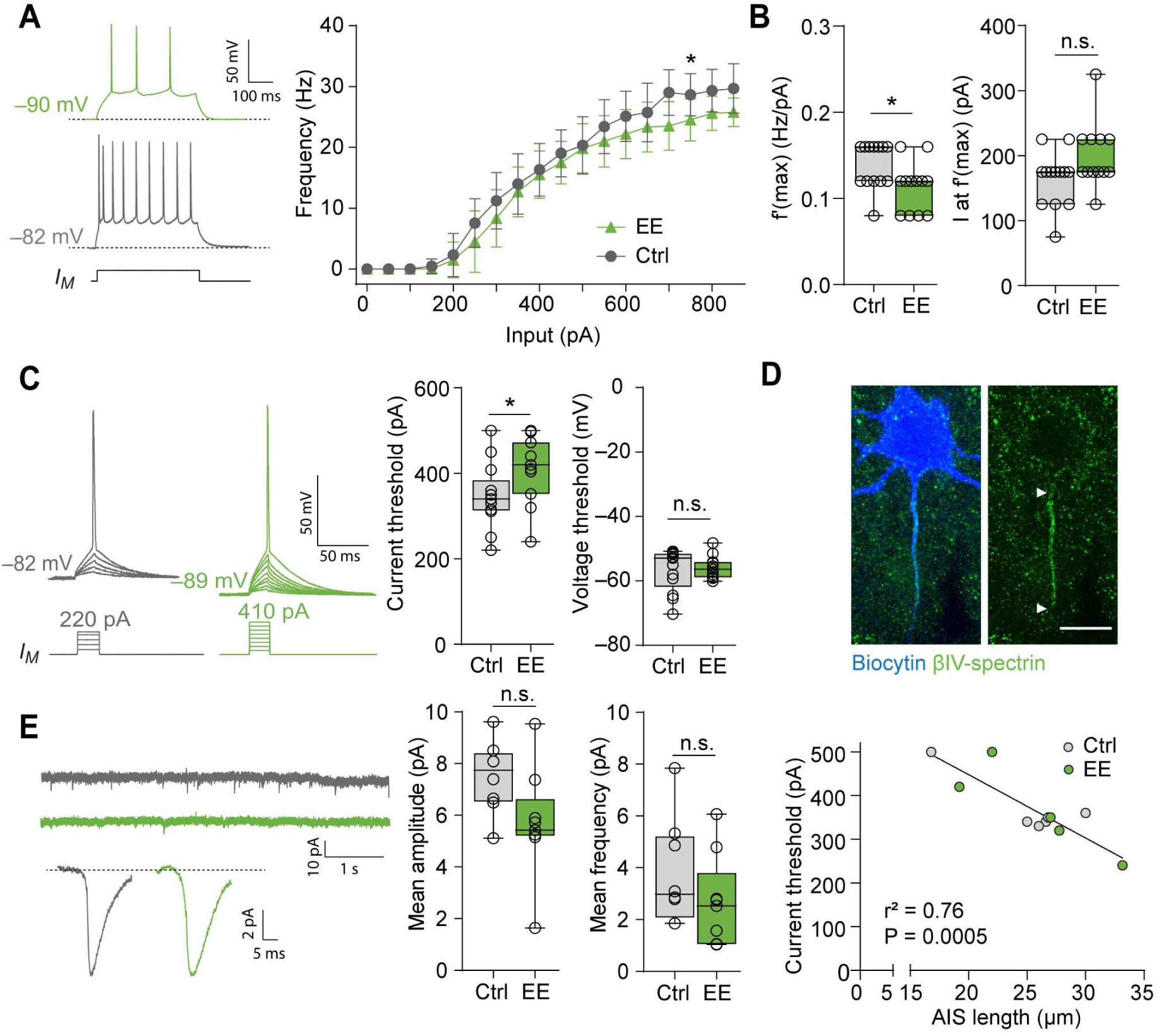
Decreased excitability in layer II/III pyramidal neurons after three hours of enriched environment (EE) **A** *Left:* Representative traces of AP trains elicited by current injection (500 ms, 250pA). Note the decreased firing frequency in the EE neuron (green trace). *Right:* Input/frequency relationship as determined by 500 ms long injections of increasing currents. The EE group showed significantly decreased firing frequencies (Two-way ANOVA (current x EE) *P* < 0.0001 for current injection and EE, *P* = 0.375 for the interaction, Sidak’s multiple comparison test \**P* = 0.035 *n* = 13 cells for Ctrl, 12 cells for EE) **B** Analysis of the maximum slope f’(max) of the I/f curve and the current at the maximum slope I at f’(max) for the respective neuron. The maximum slope was significantly lower in the EE group (unpaired *t*-test *\*P* = 0.032). The current at the maximum slope was unchanged (unpaired t-test *P* = 0.058). *n* = 13 cells for Ctrl, 12 cells for EE **C** *Left*: Representative traces of single APs elicited by 20 ms current injections. Injected current was increased in 10 pA increments to determine the current threshold. *Right*: Analysis of current and voltage threshold for threshold APs. EE neurons had significantly higher current threshold (unpaired *t*-test \**P* = 0.033). The voltage threshold was unchanged (Mann-Whitney test P = 0.73). *n* = 13 cells for Ctrl, 12 cells for EE. **D** *Top*: Representative confocal image of a neuron from the EE group that was filled with biocytin (blue) for post-hoc determination of AIS length via colabeling with βIV-spectrin (green). Arrows indicate start and end of AIS. Scale bar = 10 µm. *Bottom*: Correlation analysis of the relationship between AIS length and current threshold. Results of linear regression analysis indicated in figure (*n* = 11 cells). **E**: *Left:* Representative traces of postsynaptic currents (PSCs) recorded at −70 mV. Top trace shows 10 s of recording. Bottom trace depicts the averaged PSCs across the entire recording session of 2 min for two example cells. *Right*: Mean amplitude and frequency of PSCs. No significant difference was observed (unpaired *t*-test, amplitude: *P* = 0.0577, frequency: *P* = 0.224, *n* = 8 cells for Ctrl, *n* = 9 cells for EE). Boxplots indicate median with 25 to 75% interval and error bars show minimum to maximum values.

The above results from the relationship between c-Fos expression and AIS length suggest that it is the activity of an individual neuron that triggers homeostatic AIS scaling. To test this link more directly, we subjected acute brain slices to 1 h, 3 h, or 6 h of elevated extracellular potassium (8 mM KCl) to induce a chronic increase in activity (Fig. 6A, B). In a second alternative approach, increased activity was induced by adding 15 µM bicuculline (Bicu) to the standard ACSF (Fig. 6A, C). Strikingly, both experiments yielded a similar result: AIS length, as analyzed in layer II/III of S1Bf in acute brain slices, rapidly shortened after only 1 h of incubation in either KCl or Bicu (KCl and Bicu: One way ANOVA *P* = 0.01, Fig. 6B, C). The length changes were on average ∼8 and ∼10 µm for KCl and Bicu, respectively. AIS length then steadily increased again, reaching baseline levels after 6 h of incubation, thus mimicking the rapid reversal of AIS shortening that occurred in the *in vivo* paradigm.

**Fig. 6:**
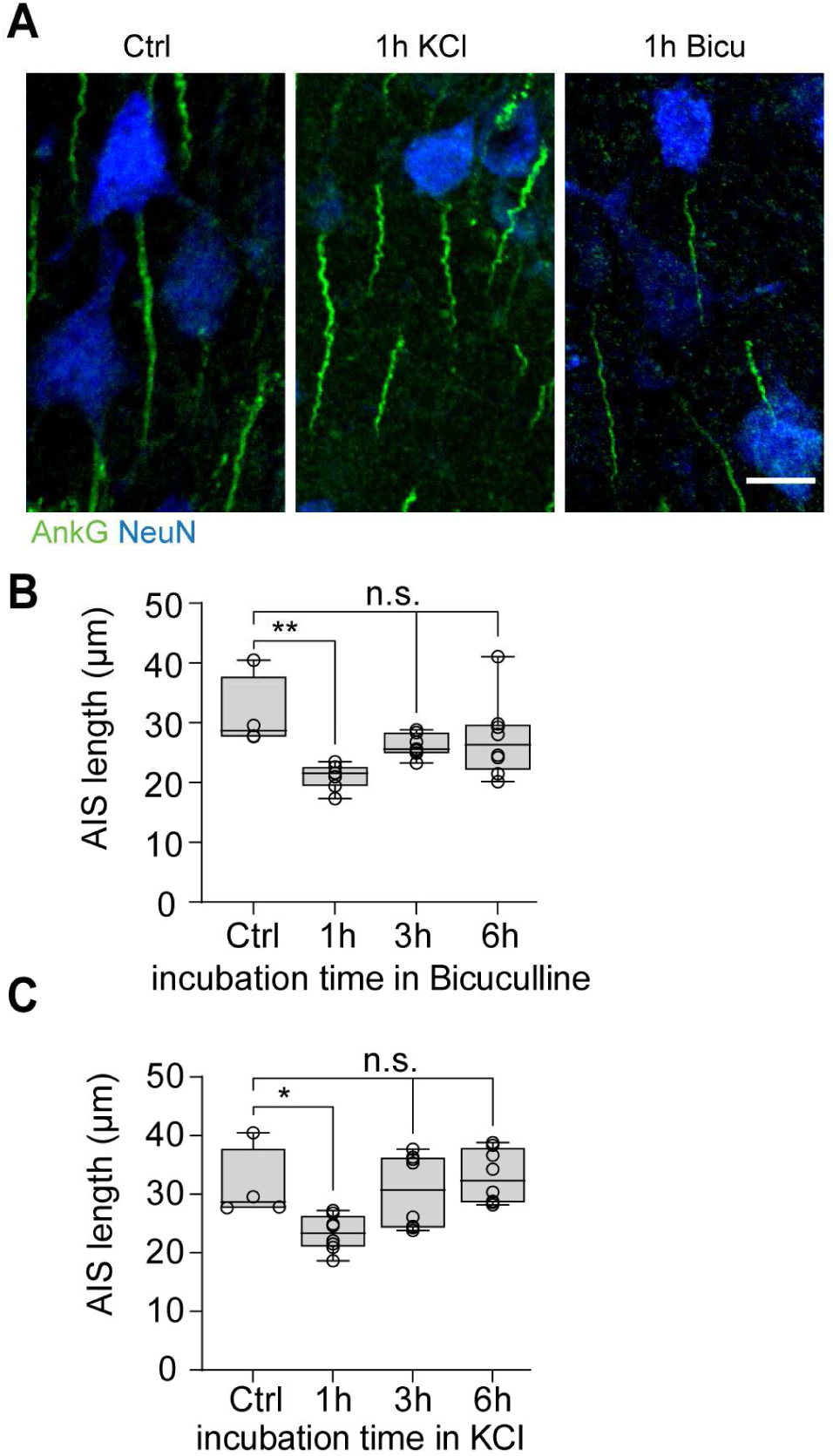
Rapid AIS length shortening in acute slices after one hour increased activity. **A:** Representative confocal images of layer II/III neurons in S1BF of acute slices that were subjected to 1 h of KCl (8 mM) or bicuculline (15 µM) treatment. Note the shorter AIS length in comparison to Ctrl conditions. Scale bar 10 µm **B**: Analysis of AIS length changes in layer II/III S1BF after exposure to bicuculline: 1 h of incubation in 15 µM Bicu leads to a rapid shortening of AIS length (One-way ANOVA *P* = 0.01, Dunnett’s multiple comparisons test \*\**P* = 0.004). AIS length returns to baseline after 3 h and 6 h of Bicu treatment. *n* = 2 mice, *n* = 3 - 4 slices per condition, > 200 AIS/condition **C**: Analysis of AIS length changes in layer II/III S1BF after exposure to elevated extracellular KCl: 1h of incubation in 8 mM KCl leads to a rapid shortening of AIS length (One-way ANOVA *P* = 0.01, Dunnett’s multiple comparisons test \**P* = 0.0346). AIS length returns to baseline after 3 h and 6 h of KCl treatment. *n* = 2 mice, *n* = 3 - 4 slices per condition, > 200 AIS/condition.

In summary, our data suggest that the increase in neuronal activity elicited by the exposure to increased sensory input led to detectable changes in AIS length and consequently homeostatic scaling of intrinsic excitability of single neurons under behaviorally relevant and physiological conditions.

## Discussion

The present study identifies AIS plasticity possibly acting as a homeostatic scaling mechanism to maintain a dynamic equilibrium for individual neurons after changes in network activity. This is supported by data showing temporally distinct, bidirectional changes of AIS length and neuronal excitability after manipulating network activity in one and the same sensory system *in vivo*.

Critical periods of cortical plasticity rely on sensory experience for maturation as has been shown for numerous sensory systems (Arakawa and Erzurumlu, 2015; Barkat et al., 2011; Espinosa and Stryker, 2012; Hensch and Fagiolini, 2005; Katz and Shatz, 1996; Kirkby et al., 2013). In this context, the rodent whisker-to-barrel system is one of the most studied sensory systems to investigate the effects of sensory experience and experience-driven modulation of network state on neuronal circuit formation, not only during development but also in the adult (reviewed in (Feldmeyer et al., 2013; Jamann et al., 2018; Petersen, 2019)). The whisker-to-barrel network is active from the emergence of whiskers onward (already during the late embryonic period), and serves important functions immediately after birth in rodents ((Sullivan et al., 2003), reviewed in (Arakawa and Erzurumlu, 2015)). At the network level, early spontaneous whisker deflections as well as passive stimulation by the mother and littermates trigger cortical bursts (Akhmetshina et al., 2016). These synchronous and spatially confined spindle bursts persist during the early postnatal period (Khazipov et al., 2004; Yang et al., 2016) and begin to wear off from P3 onwards (McCabe et al., 2007). Around P12, network activity levels are desynchronized (Golshani et al., 2009). In keeping with this developmental time line, we found that from the time of first whisker emergence prior to birth (E20) throughout the first two weeks of postnatal development, AIS in cortical layers II/III and V gradually increase in length (Fig. 1), with layer V AIS elongating before layer II/III neurons.

Around P12, mice exhibit mature rhythmic whisking behavior as indicated by adult frequencies and amplitudes (Arakawa and Erzurumlu, 2015). At the same time, information processing in S1BF changes profoundly in a layer-specific manner (van der Bourg et al., 2017). With the sudden onset of directed sensory input around P12-13, and hence an elevation of cortical network activity, we find that AIS shorten, subsequently reaching an adult length (Fig. 2). While this tri-phasic AIS maturation profile was more pronounced in visual cortex (Gutzmann et al., 2014; Schlüter et al., 2017), similar observations have been made in the current study. After the onset of active whisking and explorative behavior at P12-13 (Landers and Philip Zeigler, 2006), a reduction in AIS length was observed, followed by a gradual length increase throughout adulthood. These phasic length changes coincide with several developmental programs. For example, the second postnatal week is a period of wiring of intra- and interlayer connections in S1BF (Lendvai et al., 2000; Stern et al., 2001; Wen and Barth, 2011), which likely represents homeostatic changes. These ultimately lead to balanced network activity, especially regarding the emergences of excitation/inhibition balance in cortical networks during this time (Maravall et al., 2004; Micheva and Beaulieu, 1996). Also, GABAergic synapses originating from cortical chandelier cells (ChC) appear at the AIS between P12 – P18 in layer II/III of mouse S1 (Pan-Vasquez et al., 2020; Tai et al., 2019)). The period of AIS shortening therefore coincides with the peak of ChC synapse formation and overall increase in inhibitory conductance (Kobayashi et al., 2008; Zhang et al., 2011).

### Long-term deprivation triggers AIS plasticity and changes neuronal output parameters

We found that long-term sensory deprivation triggers AIS elongation in layer II/III, but not V (Fig. 2), rendering layer II/III neurons more excitable (Fig. 3). The lack of sensory input for the first two weeks is most likely the direct source of these changes. In support of this idea, a study characterizing intrinsic firing properties of layer II/III neurons in rat S1BF at P12, P14, and P17 found that overall excitability slightly decreased with age (Maravall et al., 2004). The authors performed whisker deprivation from P9 and found no effect on the spiking properties of these neurons, suggesting that an intrinsically mediated shift in excitability is independent of sensory input. Similarly, we found that brief deprivation (P10-15), even during the peak of important critical periods for short-range synapses (layer IV to layer II/III ipsilateral) and long-range synapses (layer II/III to layer II/III contralateral; (Sehara et al., 2010; Wen and Barth, 2011)), does not lead to significant AIS length changes (Fig. 2). Only complete deprivation from birth to P15 elicits detectable and significant structural and functional plasticity (Figs. 2, 3). Nevertheless, even though deprivation was continuous throughout the early postnatal period, developmental AIS shortening still occurred, albeit in an alleviated manner. This suggests that developmental AIS plasticity is partially mediated by sensory input, but possibly also by intrinsic, sensory-independent, developmental programs, that are genetically predetermined. Possibly, the type of non-invasive deprivation chosen in our experiments (touch sensors are left intact) affects the strength of overall AIS remodeling. This may allow for a remaining, albeit weak, general activation of the whisker-to-barrel pathway as opposed to methods of more complete elimination of sensory input (e.g. cauterization of the infraorbital nerve or whisker plucking). The passive touch information through pressure on the snout might already be sufficient to drive developmental, genetically-encoded programs.

### Layer-specific AIS plasticity

We unexpectedly also observed AIS plasticity in layer II/III of adult animals when exposed to long-term whisker deprivation (Fig. 2), which did not occur in layer V. These findings are in support of previous studies showing that supragranular layers in mouse S1BF retain plasticity in response to long-term deprivation into adult ages (Glazewski and Fox, 1996). Furthermore, studies in sensory cortices of various rodent species have outlined a layer-specific ability for plastic adaptation reflected by changes in spine dynamics, synaptic plasticity or direct stimulation responses both during development as well as at mature ages (Jiang et al., 2007; Skibinska et al., 2000; van der Bourg et al., 2017; van der Bourg et al., 2019). These studies showed that synaptic scaling and spine dynamics are a predominant feature of supragranular layers and to a much lesser extent for infragranular layers, underscoring our data showing layer-specific AIS plasticity. Overall, this might reflect different roles for supra-vs. infragranular neurons during events of developmental and behavior-induced plasticity.

### Bidirectional, temporally diverse forms of AIS plasticity

An important finding of the present study is the two distinct temporal scales at which AIS plasticity operates: sensory deprivation leads to AIS elongation and increased excitability if employed over periods of at least 2 weeks. On the other hand, AIS shortening and a decrease of excitability can be produced within an hour of sensory stimulation *in vivo* (Figs. 3, 4) or via increased neuronal activity *ex vivo* (Fig. 5). The structural changes were consistent and correlated with bidirectional changes in AP generation, supporting the idea that the anatomical changes led to functionally relevant shifts in intrinsic excitability. The two directionally and temporally distinct forms of AIS plasticity may reflect distinct underlying molecular mechanisms. Corroborating data for a long-term plasticity mechanism has been provided by studies of the avian and mammalian auditory system, where at least one week of deprivation was required in order to elicit AIS elongation and changes in neuronal excitability (Kim et al., 2019; Kuba et al., 2010). Interestingly, our data suggests that elongation only occurs at the distal end of the AIS, which requires protein biosynthesis and axonal transport of the macromolecular complexes via the microtubule network towards the distal site, and subsequent protein assembly (Freal et al., 2019). This energetically “expensive” task is presumably time-consuming, particularly considering the vast downstream network of molecular binding partners localized in the AIS (Hamdan et al., 2020).

On the other hand, rapid AIS plasticity as reported after significant elevation of neuronal activity *in vitro* seems to be driven by Ca^2+^ signaling-related mechanisms and includes proteolysis in a calpain-dependent manner, which occurs on a much faster time scale of the disassembly of AIS proteins (Benned-Jensen et al., 2016; Evans et al., 2015; Evans et al., 2013; Schafer et al., 2009). Here, we found that rapid AIS plasticity occurs after a brief increase in activity during a behaviorally-relevant context *in vivo*, with AIS length reduction already 1 h after exposure to EE (Fig. 4, 5). Similar results from acute slices exposed to KCl or bicuculline further support our hypothesis that AIS disassembly is generated by elevated neuronal activity (Fig. 6). Together with the deprivation-induced AIS length changes, the rapid AIS length reduction provides evidence of a homeostatic mechanism at the AIS to normalize activity in reaction to changes in network activity.

### Functional consequences of AIS plasticity

Interestingly, both in the long-term deprivation experiments as well as the rapid activity-induced shortening, AIS length was a strong predictor of AP current threshold (Figs. 3, 5). Whereas the AP waveform was conserved, AIS length changes appeared to selectively determine the input current to reach threshold, consistent with theoretical predictions and experimental studies observing length scaling (Kole and Brette, 2018). The input current-frequency plots in the deprivation group confirmed that excitability was increased around current threshold levels, whereas maximum firing frequencies remained unaffected. Furthermore, our data on the upregulation of c-Fos demonstrate that AIS plasticity occurs especially in those neurons that receive strong sensory activation. These results support our hypothesis that rapid AIS plasticity operates as a homeostatic scaling mechanism for activity regulation at a cell-autonomous level (Turrigiano, 2012). Our data show that after deprivation, synaptic input in layer II/III became strengthened and more frequent as indicated by the spontaneous postsynaptic currents. These results differ from previous studies showing that whisker deprivation weakens excitatory layer IV to layer II/III input, while layer IV neuronal responses remain unchanged in rat S1 (Bender et al., 2006; Glazewski and Fox, 1996). We cannot exclude cell-intrinsic changes in layer IV neurons in our model following the whisker deprivation. Further studies are required to investigate whether AIS plasticity also occurs in layer IV as well as at preceding thalamic relay stations.

In summary, our study provides evidence for a direct involvement of AIS plasticity in homeostatic scaling mechanisms in S1BF during development and behaviorally-evoked network changes within physiological contexts *in vivo*. AIS plasticity seems to be a direct response to changes in neuronal activity and occurs in an unexpectedly short timescale. Furthermore, evidence for a homeostatic role is provided by data indicating that AIS plasticity is both bidirectional and reversible. Further experiments are required to establish the capacity of single neurons to scale their AIS and being able to rescue or accelerate the plasticity responses with molecular means. Obtaining such insights and addressing these mechanistic questions would require live-imaging of single AIS *ex vivo* or *in vivo*. Once adequate live AIS reporters are available for *in vivo* application, such studies will provide unique opportunities to further our understanding of AIS plasticity in physiological contexts.

## Acknowledgments

The authors wish to thank Silke Vorwald for outstanding laboratory support and Johannes Roos for help with programming the AISuite software. Furthermore, we are indebted to Rudolph Schubert, Jana Maurer, and Martin Kaiser for providing expertise, support and technical equipment for establishing the electrophysiological experiments. We also thank Paul Jenkins and Vann Bennett for their intellectual and technical support regarding Ankyrin-G immunoblotting. We would like to thank Petra Wahle for valuable discussion and input during the writing process of the manuscript. This work was supported by the Core Facility Life Cell Imaging Mannheim (LIMA) at the MCTN (DFG-INST 91027/10-1 FUGG). The study was funded by the German Research Foundation (DFG, SFB 1134, TP A/03) to C.S. and M.E. and the Netherlands Research Council (NWO *Vici* grant 865.17.003) provided to M.H.P.K.

## Author contributions

Conceptualization, N.J. and M.E.; Methodology, N.J., R.W., J.S., and M.E.; Investigation, N.J., DD., C.C., and M.E.; Writing – Original Draft, N.J., M.H.P.K., and M.E..; Writing – Editing and Feedback, N.J., J.S., M.H.P.K., and M.E.; Visualization – N.J., M.H.P.K., and M.E.; Resources, N.J., D.D., C.C., J.S., and M.E.; Supervision, M.E.; Funding Acquisition, C.S., M.H.P.K., and M.E.

## Material and Methods

### Animals

All animal procedures were carried out in accordance with the recommendations of the Animal Research Council of the Medical Faculty Mannheim, Heidelberg University and were approved by the State of Baden-Württemberg and compliant with EU guidelines. Mice of mixed gender from the wildtype C57BL/6JRj strain (Janvier Labs, France) were maintained with food and water *ad libitum* on a regular 12 h light/dark cycle. Electrophysiological experiments were conducted between 3 and 8 h after onset of the light phase.

### Developmental and deprivation study

For the analysis of barrel cortex development, a total of 5-6 brains were analyzed in each of the following age groups: E20, P1, P3, P7, P10, P13, P15, P21, P28, P35, P45, P180. The maturation of AIS length is a robust indicator of developmental progression (Gutzmann et al., 2014; Kuba et al., 2010; Schlüter et al., 2017) and was therefore chosen as the key AIS parameter in this study. At least 100 AIS per animal were examined in S1BF in layers II/III and V, respectively. For some of the age points, an additional three animals were sacrificed for Western blot analysis of AIS scaffolding protein expression.

For sensory deprivation experiments, animals were subjected to daily bilateral whisker trimming using a curved eye scissor from P0, P10 or >P100 to different end points (see Table 1, Fig. 2A). Whisker regrowth was monitored daily under a binocular microscope and constantly kept below 1 mm of length. In some experimental groups, whiskers were allowed to regrow (see Table 1, Fig.2A). Animals older than P15 were briefly anesthetized with isoflurane prior to handling. Adult mice (> P100) were anesthetized with 40 mg Ketamine/5 mg Xylazine i.p. to minimize stress during trimming. To exclude any effects of handling or anesthesia on AIS length, control experiments were performed (see Table S3, Fig. S1D). Electrophysiological measurements were performed in 7 animals per group, yielding about 15 – 20 cells per group and analysis.

**Table 1.**
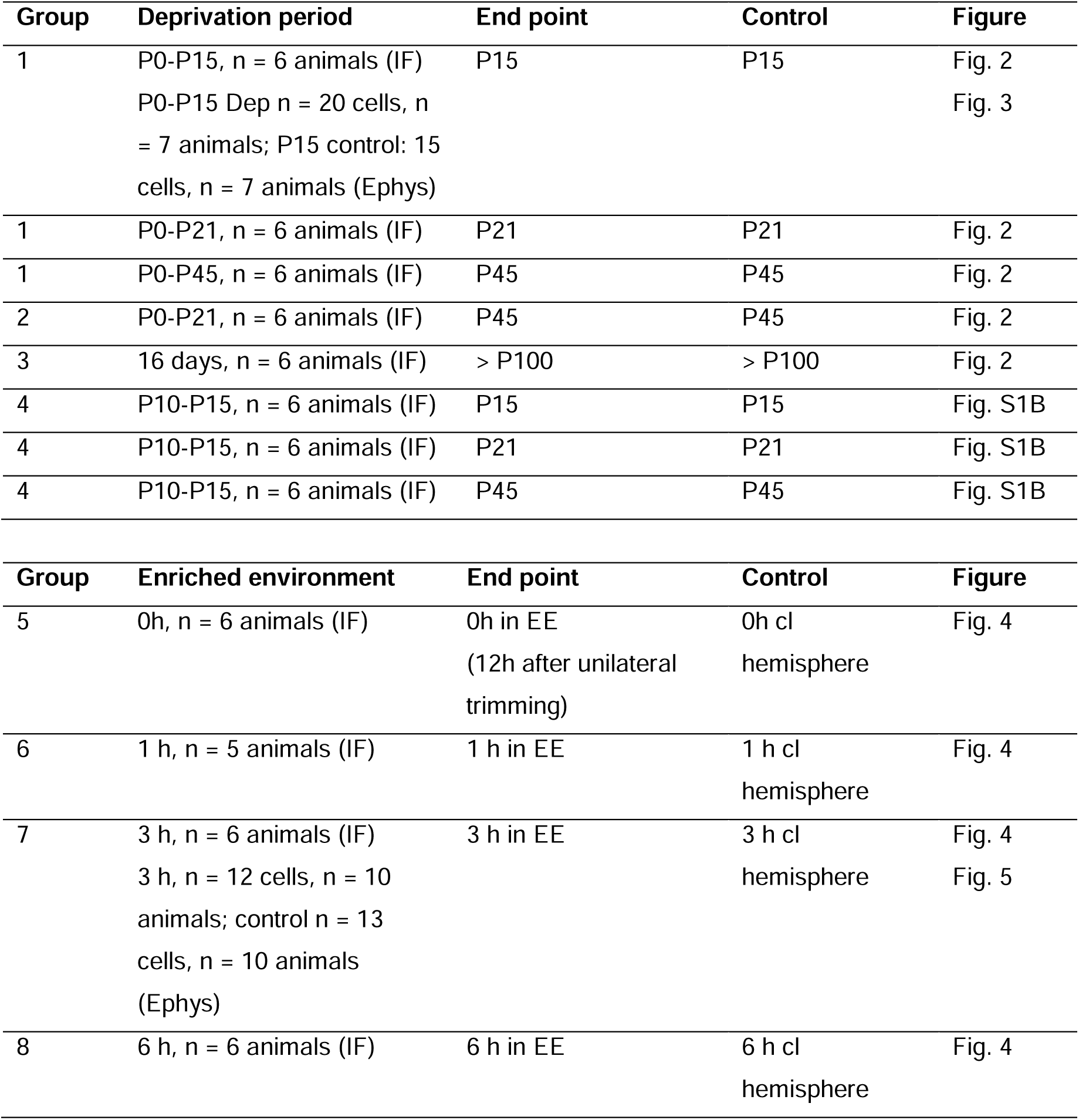
Summary of experimental groups in deprivation and enriched environment conditions. *IF* immunofluorescence, *Ephys* electrophysiological recording, *cl* contralateral

| Group | Deprivation period | End point | Control | Figure |
| --- | --- | --- | --- | --- |
| 1 | P0-P15, n = 6 animals (IF)<br>P0-P15 Dep n = 20 cells, n<br>= 7 animals; P15 control: 15<br>cells, n = 7 animals (Ephys) | P15 | P15 | Fig. 2<br>Fig. 3 |
| 1 | P0-P21, n = 6 animals (IF) | P21 | P21 | Fig. 2 |
| 1 | P0-P45, n = 6 animals (IF) | P45 | P45 | Fig. 2 |
| 2 | P0-P21, n = 6 animals (IF) | P45 | P45 | Fig. 2 |
| 3 | 16 days, n = 6 animals (IF) | > P100 | > P100 | Fig. 2 |
| 4 | P10-P15, n = 6 animals (IF) | P15 | P15 | Fig. S1B |
| 4 | P10-P15, n = 6 animals (IF) | P21 | P21 | Fig. S1B |
| 4 | P10-P15, n = 6 animals (IF) | P45 | P45 | Fig. S1B |
| Group | Enriched environment | End point | Control | Figure |
| 5 | 0h, n = 6 animals (IF) | 0h in EE<br>(12h after unilateral<br>trimming) | 0h cl<br>hemisphere | Fig. 4 |
| 6 | 1 h, n = 5 animals (IF) | 1 h in EE | 1 h cl<br>hemisphere | Fig. 4 |
| 7 | 3 h, n = 6 animals (IF)<br>3 h, n = 12 cells, n = 10<br>animals; control n = 13<br>cells, n = 10 animals<br>(Ephys) | 3 h in EE | 3 h cl<br>hemisphere | Fig. 4<br>Fig. 5 |
| 8 | 6 h, n = 6 animals (IF) | 6 h in EE | 6 h cl<br>hemisphere | Fig. 4 |

### Enriched Environment

P28 mice were whisker-trimmed unilaterally on the evening prior to the experiment so that the corresponding hemisphere was serving as an internal control (Fig. 4A). 12 h later, mice were placed in a large cage containing different types of bedding, wall textures, and various novel objects for either 0, 1, 3 or 6 h, and the cage was placed in the dark to increase explorative behavior (Fig. 4, Fig. 5). No significant changes between the two hemispheres were observed in the 0 h group. This allowed us to use the hemisphere corresponding to the trimmed whisker pad as a control for the hemisphere experiencing the increased sensory input within the same animal.

Immediately after the indicated times, animals were either subjected to cardiac perfusion for immunofluorescence or sacrificed for electrophysiological recordings (see section below and Table 1). One experimental group was placed back in a normal home cage for an additional 3 h before being sacrificed (Fig. 4E). To control for any effects of the unilateral whisker trimming, one group was trimmed and perfused the next morning without being placed in an enriched environment (0 h group, Fig. 4D). We included 5-6 animals per group for immunofluorescence and 10 per group for electrophysiology (n = 13 cells Ctrl, 11-12 cells EE)

### Immunofluorescence

For the developmental and deprivation studies, P0 - P7 animals were decapitated and brains were dissected in ice-cold 0.1 M phosphate buffer (PBS), fixed for 5 min by immersion in 4% paraformaldehyde (PFA, in 0.1 M PBS, pH 7.4) at 4°C and cryoprotected in 10% sucrose (overnight), followed by 30% sucrose (overnight) at 4°C. Animals P10 and older were exsanguinated with 0.9% NaCl under deep anesthesia with Ketamine (120 mg/kg BW)/Xylazine (16 mg/kg BW) and perfusion-fixed with ice-cold 4% PFA for five minutes. Brains were then removed from the skull and were cryoprotected in 10% sucrose (overnight) followed by 30% sucrose (overnight) at 4°C. Tissue was trimmed to a block including S1BF and embedded in Tissue Tek (Sakura Finetek). Multichannel immunofluorescence staining was performed on 20 µm sections collected directly on slides and as described previously (Gutzmann et al., 2014). Briefly, slices were incubated in blocking buffer (1% BSA, 0.2% fish skin gelatine, 0.1% Triton in 0.1 M PBS) for at least 60 min and subsequently incubated in primary antibodies overnight at 4°C. After washing, slices were incubated for at least 90 min in secondary antibodies in the dark at room temperature. For preservation of immunofluorescence, slices were mounted in a mounting medium with anti-fading effect (Roti-Mount FluorCare, Carl Roth). After omission of the primary antibodies and application of only secondary antibodies, no specific immunolabeling was observed. Antibodies were further tested for validity as outlined in Table 2. Acute slices from electrophysiological recordings were fixed in 4% PFA for 20 min. Staining was carried out as described above, however with prolonged incubation periods to allow for sufficient penetration of the tissue by the antibody. We frequently observed that in biocytin-filled neurons, AIS stainings appeared weaker. Consequently, only AIS with clearly detectable start and end points were chosen for the length correlation analysis (Fig. 3D and 5D).

**Table 2.**
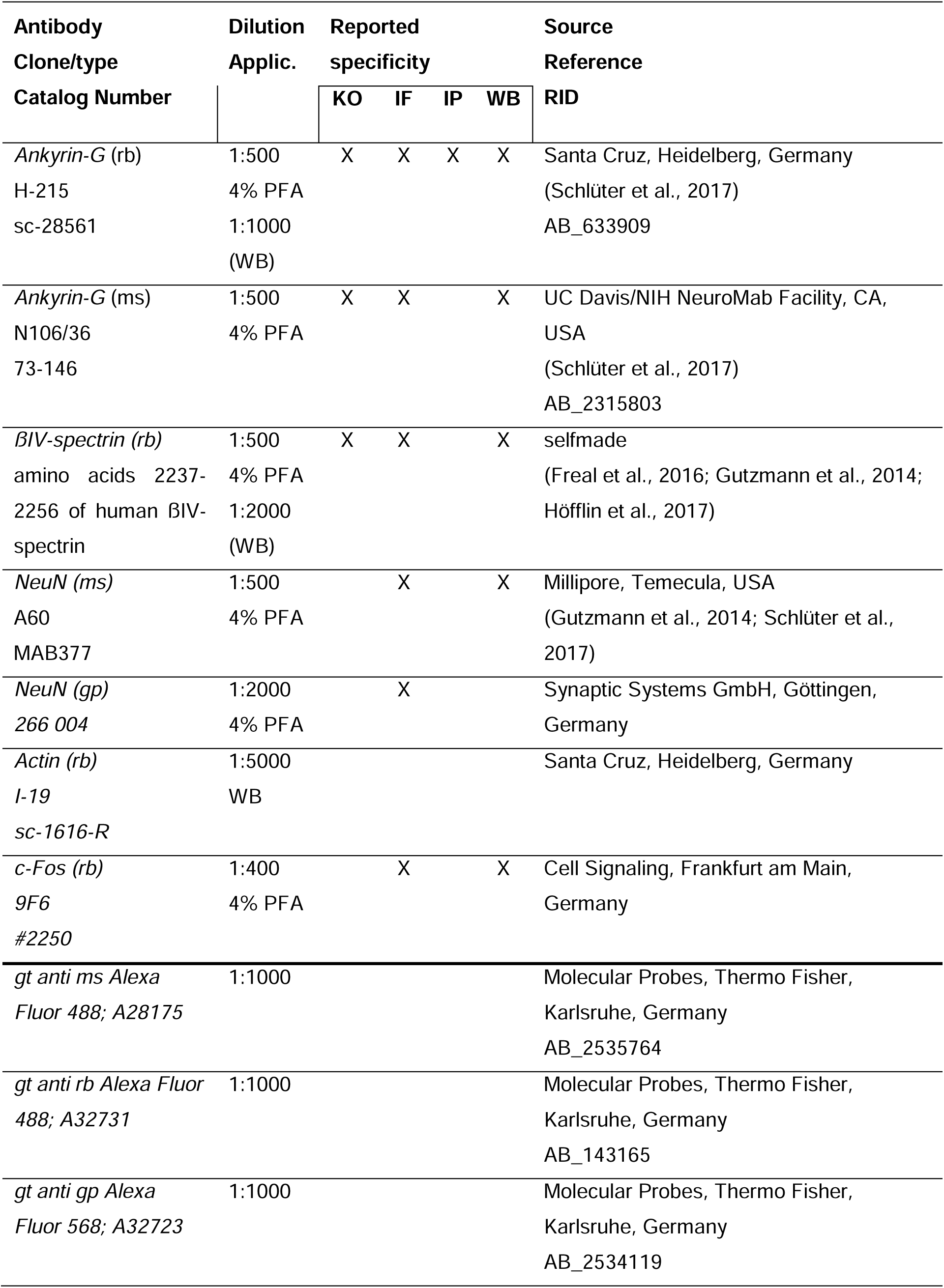
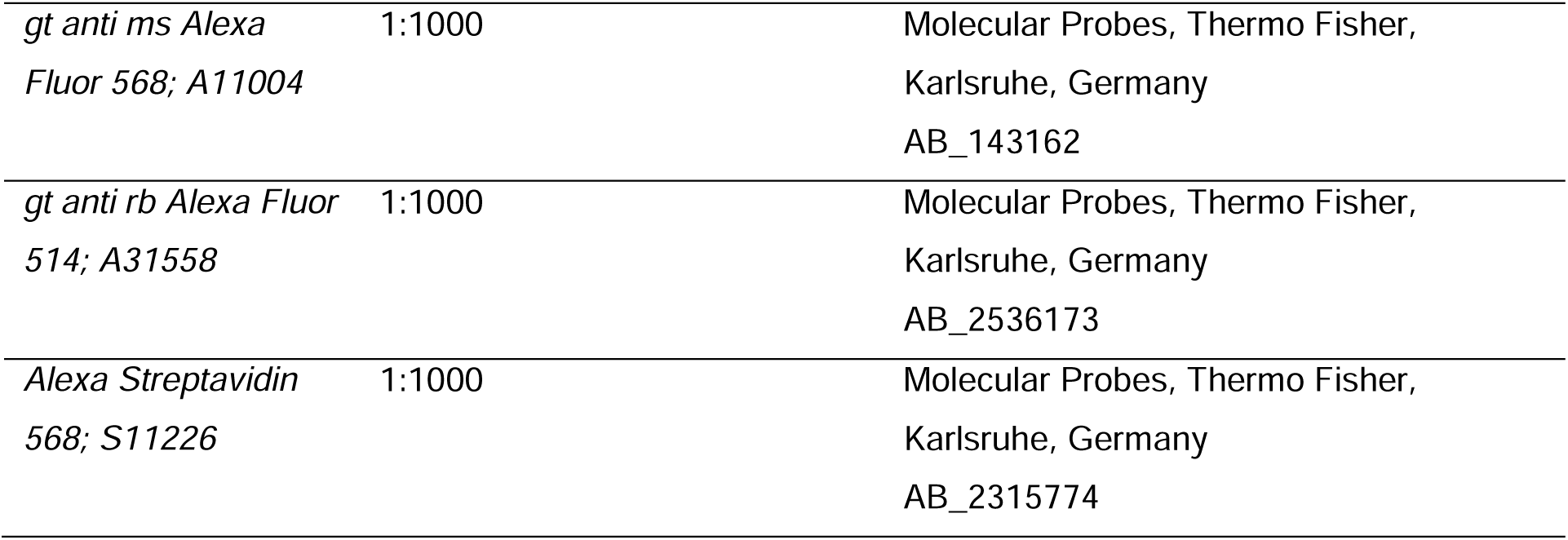
Specification of antibodies (catalog number, working dilution, fixation of tissue, previously conducted controls, sources and references where available). *KO* absence of immunostainings in knock out animals, *IF* immunofluorescence, *IP* immunoprecipitation, *WB* western blot, *rb* rabbit, *ms* mouse, *gp* guinea pig

| Antibody<br>Clone/type<br>Catalog Number | Dilution<br>Applic. | Reported<br>specificity |  |  |  | Source<br>Reference<br>RID |
| --- | --- | --- | --- | --- | --- | --- |
|  |  | KO | IF | IP | WB |  |
| *Ankyrin-G* (rb)<br>H-215<br>sc-28561 | 1:500<br>4% PFA<br>1:1000<br>(WB) | X | X | X | X | Santa Cruz, Heidelberg, Germany<br>(Schlüter et al., 2017)<br>AB_633909 |
| *Ankyrin-G* (ms)<br>N106/36<br>73-146 | 1:500<br>4% PFA | X | X |  | X | UC Davis/NIH NeuroMab Facility, CA, USA<br>(Schlüter et al., 2017)<br>AB_2315803 |
| *βIV-spectrin* (rb)<br>amino acids 2237-2256 of human βIV-spectrin | 1:500<br>4% PFA<br>1:2000<br>(WB) | X | X |  | X | selfmade<br>(Freal et al., 2016; Gutzmann et al., 2014;<br>Höfflin et al., 2017) |
| *NeuN* (ms)<br>A60<br>MAB377 | 1:500<br>4% PFA |  | X |  | X | Millipore, Temecula, USA<br>(Gutzmann et al., 2014; Schlüter et al., 2017) |
| *NeuN* (gp)<br>266 004 | 1:2000<br>4% PFA |  | X |  |  | Synaptic Systems GmbH, Göttingen, Germany |
| *Actin* (rb)<br>I-19<br>sc-1616-R | 1:5000<br>WB |  |  |  |  | Santa Cruz, Heidelberg, Germany |
| *c-Fos* (rb)<br>9F6<br>#2250 | 1:400<br>4% PFA |  | X |  | X | Cell Signaling, Frankfurt am Main, Germany |
| *gt anti ms Alexa*<br>*Fluor 488; A28175* | 1:1000 |  |  |  |  | Molecular Probes, Thermo Fisher, Karlsruhe, Germany<br>AB_2535764 |
| *gt anti rb Alexa Fluor*<br>*488; A32731* | 1:1000 |  |  |  |  | Molecular Probes, Thermo Fisher, Karlsruhe, Germany<br>AB_143165 |
| *gt anti gp Alexa*<br>*Fluor 568; A32723* | 1:1000 |  |  |  |  | Molecular Probes, Thermo Fisher, Karlsruhe, Germany<br>AB_2534119 |

|  |  |  |
| --- | --- | --- |
| *gt anti ms Alexa Fluor 568; A11004* | 1:1000 | Molecular Probes, Thermo Fisher,<br>Karlsruhe, Germany<br>AB_143162 |
| *gt anti rb Alexa Fluor 514; A31558* | 1:1000 | Molecular Probes, Thermo Fisher,<br>Karlsruhe, Germany<br>AB_2536173 |
| *Alexa Streptavidin 568; S11226* | 1:1000 | Molecular Probes, Thermo Fisher,<br>Karlsruhe, Germany<br>AB_2315774 |

### Image acquisition and analysis

Confocal analysis was carried out on a C2 Nikon confocal microscope (Nikon Instruments, laser lines: 642, 543, and 488 nm) with a 60x objective (oil immersion, NA 1.4) and a SP5 confocal microscope (Leica, Mannheim; laser lines: 633, 561, 514, and 488 nm) with a 63x objective (oil immersion, NA 1.4). To increase the number of in-focus immunoreactive structures, stacks of images were merged for analysis (maximum intensity projection). For visualization of representative images in figures, brightness and contrast were optimized (ImageJ). Thickness of single optical sections was 0.5 µm in stacks of 10 - 20 µm total depth for cryosections and up to 40 µm in acute slices. Confocal *x*-*y*-resolution was set to and kept at 0.21 µm per pixel. Images for qualitative analysis were evaluated and enhanced for brightness and contrast in Photoshop (Adobe Systems) and FIJI (ImageJ), respectively. AIS length was measured using a self-written macro (Gutzmann et al., 2014; Höfflin et al., 2017) as well as the morphometrical software AISuite (github.com/jhnnsrs/aisuite2). This tool extend the well-established and widely used method of defining AIS start and end points as points where a predefined fluorescence threshold (relative to the maximum fluorescence intensity along a line drawn over an individual AIS) is surpassed (Grubb and Burrone, 2010). The threshold was adjusted depending on the individual staining quality and ranged from 10 - 30% of maximum fluorescence intensity. Both analysis tools were tested for inter-method reliability, revealing robust consistency of results.

### Western Blot

For mice from E20.5 - P3, processed samples included the entire cortex. For older animals, all brains were cut into 1 mm slices using a tissue matrix slicer (Zivic instruments) and single sections were visualized using a binoscope to carefully dissect only S1BF for sample preparation. Additionally, animals older than P10 were perfused transcardially with ice cold 0.9% NaCl as outlined above before removing the brain. Samples were diluted in a homogenization buffer (20 mM Tris, 0.5 M NaCl, 8 mM CHAPS, 6.4 mM EDTA, pH 7.5) containing phosphatase and protease inhibitors (Sigma-Aldrich). Samples were then homogenized via ultrasonication, lyzed for 60 min and centrifuged at 13,000 rpm for 45 min at −4°C. Protein quantification via a Bradford assay was performed and 20 µg samples containing Laemmli-buffer (2% SDS, 60 mM tris-Cl, 10% glycerol, 5% ß-mercaptoethanol, 0.01% bromphenol blue) were heated for 10 min at 70°C. Gradient gels (3% - 8% Tris-acetate protein gels, Thermo Fisher Scientific) were loaded with the lysates and run for 55 min at 150 V in Tris-tricine buffer (50 mM Tris, 50 mM tricine, 0.1% SDS). The blotting process was carried out in three steps according to previously published protocols (Engelhardt et al., 2018). This ensured transfer of both large and small proteins on the same membrane so ankG isoforms (ranging from 190 to 480kDa) as well as the loading control actin (50 kDa) could be transferred and visualized for later analysis. Blotting was performed at 550 pA in a tris-glycine buffer (25 mM tris-Base, 192 mM glycine) under constant cooling. Solutions contained 20% methanol for 30 min, 15% methanol and 0.05% SDS for 30 min, and only 0.1% SDS for another 90 min. Between each step, membrane strips already containing smaller proteins were removed. Membranes were blocked for 60 min in PBST (protein free). Primary antibodies were incubated overnight at 4°C (summarized in Table 2) and secondary antibodies were incubated for 90 min at room temperature. Protein signal was revealed with an ECL Kit (Western Bright ECl HRP substrate, Advansta) and imaged (Fusion solo, Vilber Lourmat). Analysis was carried out with Image J software. Samples were normalized against the internal loading control (actin) as well as, when comparing several gels, against a standardized sample with a consistent protein concentration run on each gel.

### Electrophysiology

In accordance with the guidelines of 3Rs, and to enable the use of several animals per litter, animal age for Fig. 3 (Group 1) ranged from P13 to P16 (both for Dep and Ctrl) and animal age for Fig. 5 ranged from P28 to P31 (EE). Mice were briefly anesthetized with isoflurane (3%) and decapitated. The brain was quickly removed and placed in ice cold sucrose-based cutting solution (206 mM sucrose, 2.5 mM KCl, 1.25 mM NaH_2_P0_4_, 25 mM NaHCO_3_, 25 mM Glucose, 3 mM MgCl, 1 mM CaCl_2_, pH 7.4), which was saturated with carbogen (95% O_2_, 5% CO_2_). 300 µm thick coronal sections containing S1BF were cut with a vibratome (VT 1200 S, Leica Biosystems). Acute slices were transferred to artificial cerebrospinal fluid (ACSF; 125 mM NaCl, 2.5 mM KCl, 1.25 mM NaH_2_PO_4_, 25 mM NaHCO_3_, 1 mM MgCl_2_, 2 mM CaCl_2_, 25 mM glucose, pH 7.4, oxygen-saturated with 95% O_2_, 5% CO_2_) and allowed to rest at room temperature for at least 15 min (EE conditions) and 30 min (deprivation conditions) before recordings began. All recordings were carried out at room temperature. For EE experiments, time after slice preparation was kept to a minimum and was monitored to exclude any rapid reversal of AIS shortening during incubation of acute slices (Fig. S4C). Slices were imaged with an upright Nikon Eclipse FN1 equipped with a DIC contrast filter. Layer II/III pyramidal neurons were visually identified and neuron type was confirmed post-hoc by the firing pattern and immunofluorescence against βIV-spectrin and the biocytin fill. Pipettes were pulled from borosilicate glass (outer diameter 1.5 mm, inner diameter 0.8 mm, Science Products) to a tip resistance of 3.5 - 5.5 MΩ and filled with intracellular solution (140 mM K-gluconate, 3 mM KCl, 4 mM NaCl, 10 mM HEPES, 0.2 mM EGTA, 2 mM Mg-ATP, 0.1 mM Na_2_-GTP), containing 3 mg/ml biocytin. Patch-clamp recordings were made with a HEKA EPC10 USB amplifier controlled by Patchmaster Software (HEKA Electronics). Signals were filtered with a 10 kHz (Filter 1) and 2.9 kHz (Filter 2) Bessel filter, digitized, and sampled at 50 kHz. The liquid junction potential was calculated to be –12 mV and corrected for post-hoc. Fast and slow capacitances were compensated for in cell-attached and whole-cell configuration, respectively. Series resistance (*R*_s_) was constantly monitored with a –10 mV step in voltage clamp. Cells with *R*_s_ exceeding 30 MΩ during recordings were excluded from analysis. Input resistance (*R*_N_) was calculated from the slope of the current/voltage relationship curve from a current clamp step protocol. Resting membrane potential (RMP) was measured directly upon entering whole-cell configuration in current clamp at *I* = 0. AP properties were measured with a step protocol of 20 ms pulses increasing in 10 pA increments, starting from a holding current of *I* = 0. For firing pattern analysis (including the *I*-*f* curves), 500-ms long pulses incrementing in 50 pA steps were used to trigger AP trains. Maximum slope of the curve as well as the current at maximum slope were calculated for each group. Spontaneous postsynaptic currents (PSCs) were recorded at –70 mV for 2 min. Analyses were carried out offline with either FitMaster Software (HEKA Electronics) or OriginPro 8 (Origin lab Corporation). Current threshold was defined as the current at the first 20 ms pulse that reliably elicited an AP. Voltage threshold was determined as the point where the first time derivative exceeded 50 V s^−1^. AP amplitude was measured from voltage threshold to the AP peak voltage. AP half-width was defined as the width at the middle voltage of the rising phase between AP threshold and peak. For phase plot analysis, the first temporal derivative (V s^−1^) was plotted against the voltage (V), and the value at the first (AIS) and second (somatic) peak of the AP were extracted for analysis. EPSPs were detected with the automatic event detection function of AxoGraph X (AxoGraph Scientific), and mean amplitudes and frequency were calculated for each neuron.

### Statistical Analysis

Mean values and standard deviation (SD) of AIS length were calculated, plotted and analyzed in Sigma Plot 12.5 Software (Systat Software GmbH) or GraphPad Prism 8 software (GraphPad Software, Inc.). Unpaired *t*-test and Mann-Whitney test were carried out for parametric and non-parametric comparison of only two groups, respectively. Two-way ANOVA followed by appropriate post-hoc correction was applied when comparing two or more groups over several time points (details are given in the respective figure legends). In all graphs, box plots indicate the median (middle line) across all animals with min and max value (whiskers) and 25 and 75 percentiles (bottom and top border of box). P-values and number of samples are stated in each figure legend.

## Supplemental Tables

**Table S1.** Passive properties of recorded cells from deprivation (Dep) and control (Ctrl) groups

|  | P15 Ctrl (*n* = 15) | P15 Dep (*n* = 17) | P |
| --- | --- | --- | --- |
| RMP (mV) | $-81.76 \pm 3.79$ | $-78.62 \pm 5.13$ | 0.056 |
| $R_N$ (MΩ) | $303.12 \pm 73.98$ | $347.45 \pm 72.02$ | 0.092 |
| $R_S$ (MΩ) | $14.88 \pm 2.62$ | $14.73 \pm 5.04$ | 0.922 |

RMP = resting membrane potential, $R_N$ = Input resistance, $R_S$ = Series resistance. Data are indicated as mean ± SD.

**Table S2.** Passive properties of recorded cells from enriched environment (EE) and control (Ctrl) groups

|  | P15 Ctrl (*n* = 15) | P15 EE (*n* = 17) | P |
| --- | --- | --- | --- |
| RMP (mV) | $-86.59 \pm 5.53$ | $-88.56 \pm 5.70$ | 0.538 |
| $R_N$ (MΩ) | $130.92 \pm 38.07$ | $114.08 \pm 22.97$ | 0.198 |
| $R_S$ (MΩ) | $21.81 \pm 6.87$ | $21.73 \pm 5.86$ | 0.977 |

RMP = resting membrane potential, $R_N$ = Input resistance, $R_S$ = Series resistance. Data are indicated as mean ± SD.

**Table S3.**
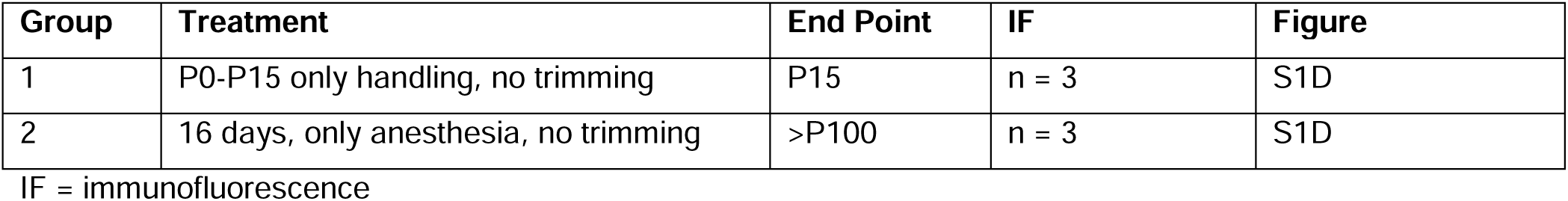
Summary of additional control groups for deprivation experiments

| Group | Treatment | End Point | IF | Figure |
| --- | --- | --- | --- | --- |
| 1 | P0-P15 only handling, no trimming | P15 | *n* = 3 | S1D |
| 2 | 16 days, only anesthesia, no trimming | >P100 | *n* = 3 | S1D |

IF = immunofluorescence

