## Supplements for "Sensory input drives rapid homeostatic scaling of the axon initial segment in mouse barrel cortex"

**Supplementary Items**

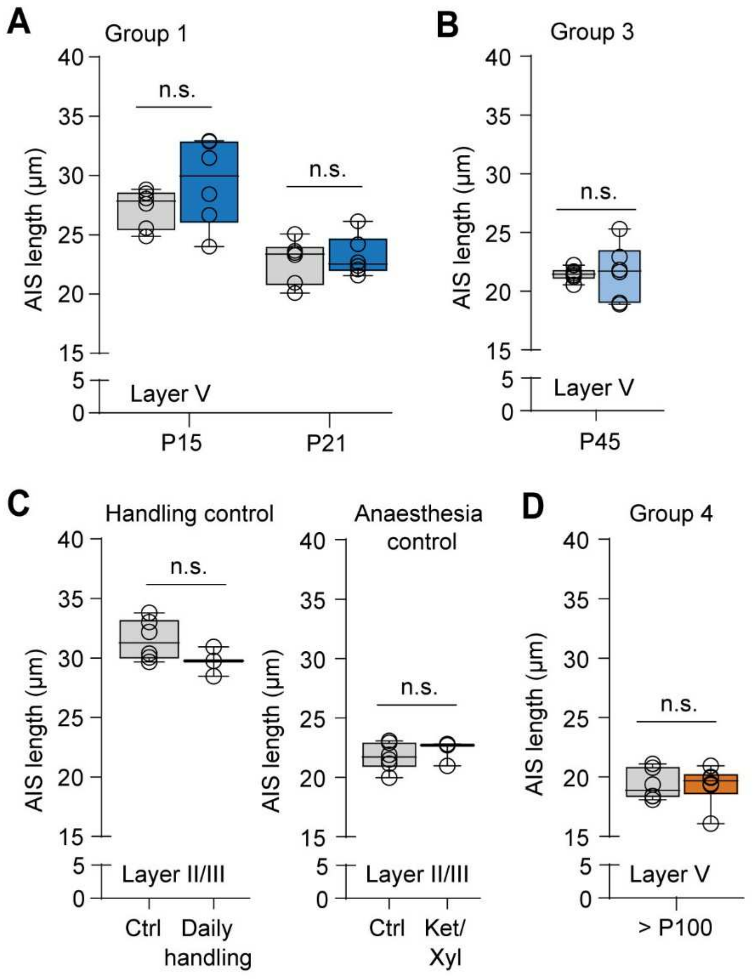


**Figure S1. Related to Figure 1: Layer V AIS do not show activity-dependent plasticity**

**A** Analysis of AIS length changes in layer V after long-term deprivation in group 1. At P15 and P21 no significant length changes were observed (Two-way ANOVA $P = 0.197$ for deprivation, $P < 0.0001$ for age, $P = 0.380$ for the interaction, Sidak's multiple comparisons $P > 0.05$, n = 6)

**B** Analysis of AIS length changes in layer V after long-term deprivation in group 3. At P45 no significant length changes were observed (unpaired t-test $P = 0.879$, n = 6)

**C** Control experiments for whisker trimming. *Left*: Equivalent to group 1, pups were handled daily from P0 to P15 but instead of trimming, whiskers were only slightly ruffled. No significant length changes were observed in this group (unpaired t-test $P = 0.1558$, n = 3). *Right*: As a anesthesia control for group 4, adult mice were given a daily dose of ketamine/xylazin for anesthesia, however whiskers were not trimmed. In the anesthetized animals no AIS length changes were observed as compared to adult controls (unpaired t-test $P = 0.625$, n = 3).

**D** Analysis of AIS length changes in layer V after long-term deprivation in group 4. In adult mice, no significant length changes were observed (unpaired t-test $P = 0.95$, n = 6)

**A – D** Boxplots indicate median with 25 to 75% interval and error bars show min. to max. values.

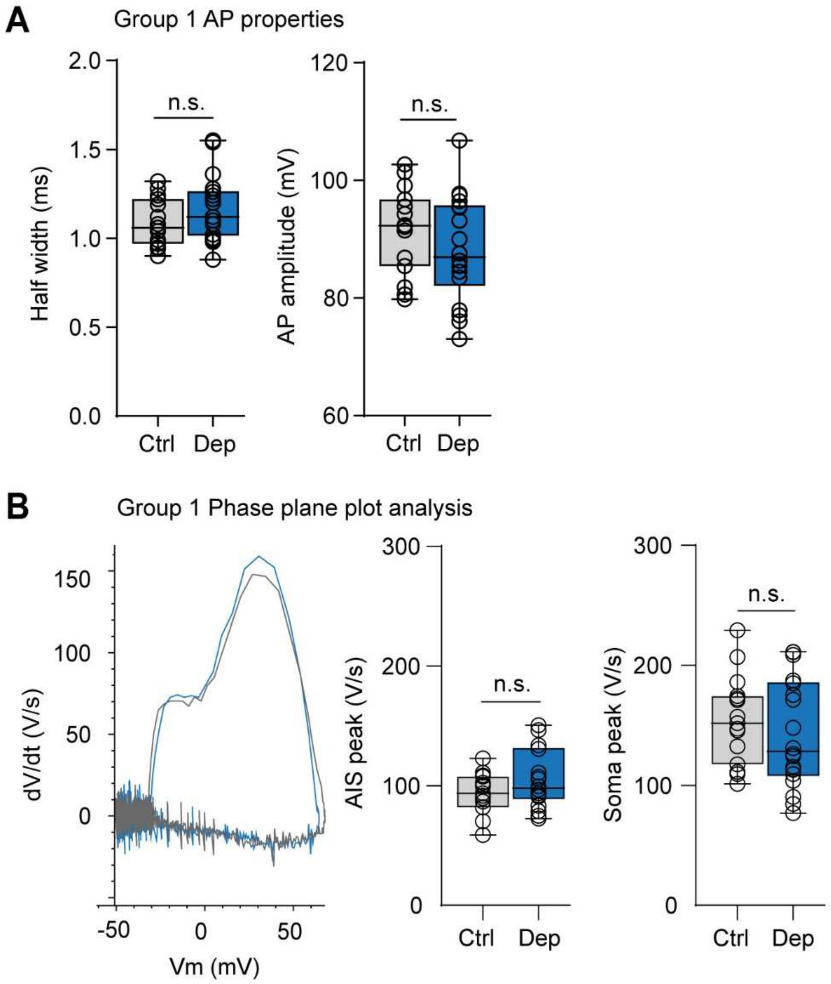


**Figure S2. Related to Figure 3: AP waveform is conserved after sensory deprivation**

**A** AP half width and AP Amplitude were not changed in group 1 after deprivation from P0 – P15 (unpaired t-test half width *P* = 0.201, AP amplitude P = 0.276, n = 15 cells for Ctrl, 20 cells for Dep).

**B**: Phase plane plot analysis of Dep vs Ctrl APs. *Left*: Representative phase plane plots of a Ctrl and Dep neuron demonstrate the similarity in AP shape. *Right*: Analysis of the first and second peak (AIS and soma peak respectively) of the phase plane plot reveals no significant difference between deprivation and control neurons (unpaired t-test AIS peak *P* = 0.106, soma peak *P* = 0.443, n = 18 cells Dep, n = 15 cells Ctrl)

**A, B** Boxplots indicate median with 25 to 75% interval and error bars show min. to max. values.

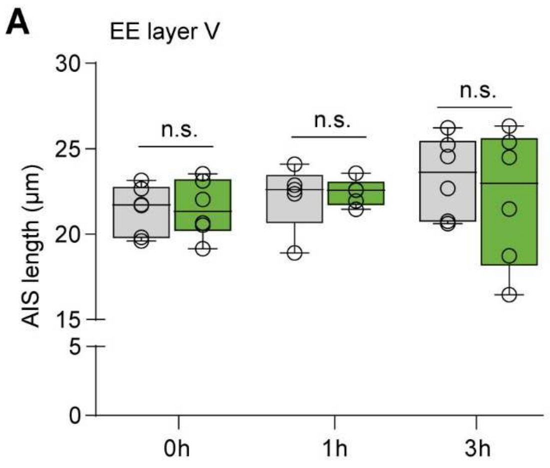


**Figure S3. Related to Figure 4: Layer V AIS length remains unaltered after exposure to an enriched environment**

Analysis of AIS length changes after EE exposure in layer V. No significant length changes were observed after 1h and 3h of EE (Two-way ANOVA $P = 0.713$ for EE, $P = 0.40$ for time, $P = 0.715$ for the interaction, Sidak's multiple comparisons $P > 0.05$ for all comparisons, $n = 5 - 6$) Boxplots indicate median with 25 to 75% interval and error bars show min. to max. values.

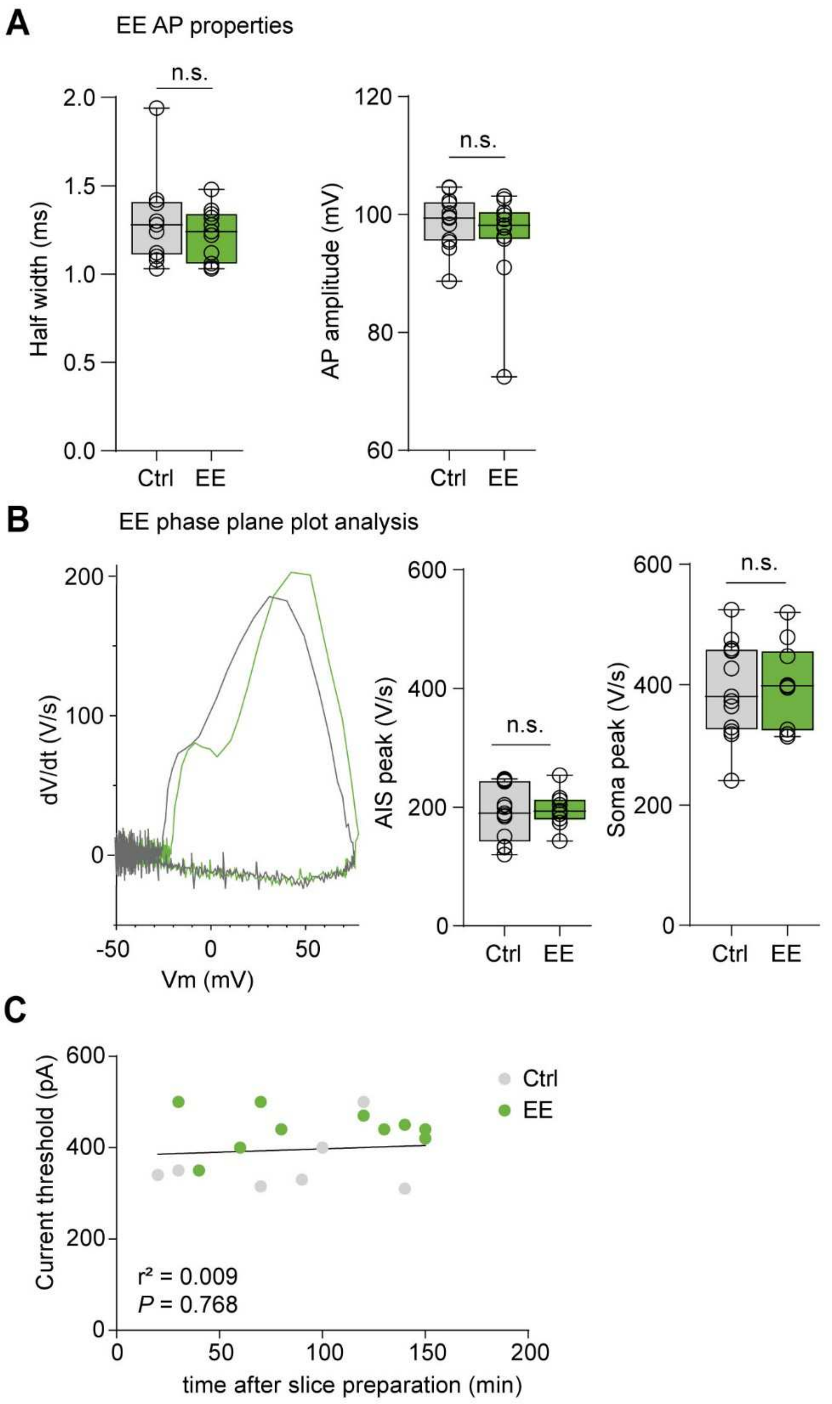


**Figure S4. Related to Figure 5: AP waveform is conserved after exposure to an enriched environment**

**A** AP half width and AP Amplitude were not changed after 3 h of EE (half width: Mann-Whitney test $P$ = 0.578, AP amplitude: unpaired t-test P = 0.34, n = 13 cells for Ctrl, n = 11 cells for EE)

**B**: Phase plane plot analysis of EE vs Ctrl APs. *Left*: Representative phase plane plots of a Ctrl and EE neuron demonstrate the similarity in AP shape. *Right*: Analysis of the first and second peak (AIS and soma peak respectively) of the phase plane plot reveals no significant difference between EE and control neurons (unpaired t-test AIS peak $P = 0.783$, soma peak $P = 0.875$, n = 10 cells EE, n = 13 cells Ctrl).

**C:** Correlation analysis of the relationship between the time after slicing and the current threshold to control for reversibility of EE effect over time during slice incubation. The results of the linear regression analysis, as indicated in the figure, reveal no correlation between time after slice preparation and current threshold.

**A, B** Boxplots indicate median with 25 to 75% interval and error bars show min. to max. values.
